# MicroRNA166-REVOLUTA-auxin module affects tuber shape, color and productivity in potato

**DOI:** 10.1101/2025.11.20.685984

**Authors:** Nikita Sunil Patil, Arati Vasav, Bhavani Natarajan, Gourav Arora, Jyoti Kumari, Anjan K. Banerjee

## Abstract

Tuber shape and size are important traits in potato. Although several factors governing tuberization have been well studied, the molecular mechanisms regulating the tuber shape and size are still elusive. Here, we demonstrate that suppression of *miR166*, which targets class III HD-ZIP transcription factors, alters tuber morphology and productivity in potato. Target mimicry of *miR166* (MIM166) in *Solanum tuberosum* ssp. *Andigena* resulted in elongated, pigmented tubers with reduced yield under short-day conditions. Transcriptome profiling of the tuberizing stolons - swollen head *vs*. stalk revealed differential expression of auxin-, cytokinin-, and gibberellin-associated genes, consistent with altered hormone levels. The coloured tubers from MIM166 line exhibited differential accumulation of cyanidin 3-glucoside, pelargonidin 3-glucoside, and delphinidin 3-glucoside. Tuber productivity in these lines was reduced possibly due to the decreased expression of *SWEET11b* and distorted vascular structures. Further, the *miR166* target-*StREVOLUTA* exhibited dynamic expression during stolon-to-tuber transition, and could modulate *StYUCCA7* expression, an auxin biosynthesis gene. Increased pigmentation and auxin accumulation in MIM166, reduced expression of auxin biosynthesis genes in REVOLUTA antisense, and low tuber yield collectively suggest *miR166*-REV as a regulatory module of auxin homeostasis in differentiating stolons that influences tuber morphology. These results reveal a previously unrecognized miRNA-mediated pathway governing storage organ shape, and extends the functional scope of the *miR166*-HD-ZIP III module beyond organ polarity.

**Significance statement:** Limited studies describe the molecular mechanisms regulating tuber shape in storage organ development. Here, we show that the *miR166-*REVOLUTA-auxin network has significant regulatory roles in potato development, influencing tuber shape, pigmentation, and productivity. These findings provide insights into molecular control of tuber morphology, uncover hitherto unknown functions of the *miR166*-REVOLUTA module in tuber crops, and broadens our understanding of *miR166*-mediated developmental regulation in plants.

## Introduction

Potato is one of the most important food crops in the world. The shape and size of potato tuber are key agronomic traits determining the tuber appearance and tuber yield. In plants, organ shape and size is determined by several regulatory genes and hormone signaling networks that affect cell division, cell expansion, and anisotropic growth (Seymour *et al*., 2013; Eldridge *et al*., 2016; Van der Knaap *et al*., 2018; Bu *et al*., 2025). In potato, a belowground modified stem-stolon undergoes developmental transition to form a tuber. Although the genetic and hormonal networks controlling stolon-to-tuber transitions have been extensively studied (Zierer *et al*., 2021), our understanding of the molecular mechanisms determining tuber shape in potato and other storage crops remains limited.

Studies in tomato and rice provided detailed insights into the molecular mechanisms regulating organ shape in fruit and grain development. In tomato, the *OVATE* locus on chromosome 2 controls the transition from round to pear-shaped fruits (Liu *et al*., 2002), while the *SUN*, *SOV1*, and *fs8.1* loci influence fruit elongation through alterations in cell number, and modulation of auxin-related genes (Wang *et al*., 2019). In potato, suppression of an Ovate Family Protein, *StOFP20* (Soltu.DM.09G010420) caused a shift from round to oval tubers. StOFP20 interacts with TONNEAU1 Recruiting Motif proteins, which in turn associate with TON2, potentially affecting cell division and expansion (Ju *et al*., 2023). In rice, grain size is determined by cell number and expansion. Genes such as *GRAIN SIZE 3*, *OsPPKL1*, *OsSPL16*, *GS5*, and the *OsMKK4-OsMPK6* regulate cell proliferation (Fan *et al*., 2006; Mao *et al*., 2010; Hu *et al*., 2012), while *SPL13*, *OsGRF4*, *OsGIF1/2/3*, and *GW7/GL7/SLG7* control cell expansion (Hu *et al*., 2015; Si *et al*., 2016; Che *et al*., 2016; Li *et al*., 2016). In potato, linkage and mapping studies identified significant quantitative trait locus (QTL) for tuber shape on chromosomes 4, 7, and 10 (Park *et al*., 2021). In a study by Fan *et al*. (2022), the *Round* (*Ro*) locus controlling potato tuber shape was fine-mapped on chromosome 10, and five candidate genes potentially responsible for determining round *versus* elongated tuber were identified. In a diploid Andean potato population, candidate genes such as BEL1-like, α-expansin, pectin biosynthesis/modification genes, SCARECROW, and ERF5 were mapped to the QTL on chromosome 10 (Lindqvist-Kreuze *et al*., 2015). Despite multiple studies identifying the QTLs and candidate genes associated with the tuber morphology, their molecular characterization is awaited.

Phytohomones, particularly auxin and gibberellin are shown to influence fruit shape in plants. Early work by Stembridge and Morrell (1972) demonstrated that gibberellins (GA_4_, GA_7_) and cytokinin (6-BAP) increased the length-to-diameter ratio in apples, with combined application being most effective. In tomato, GA_3_ application promotes fruit elongation through upregulation of cell elongation genes, whereas inhibition of gibberellin biosynthesis reduces elongation and accelerates ripening (Chen *et al*., 2020). Similarly, the *SUN* locus influences the fruit shape in tomato by modulating auxin biosynthesis, transport, and signaling genes, indicating an interplay between phytohormones and genetic regulators in organ morphology (Wang *et al*., 2019). Thus, organ shape in plants is governed by a coordinated function of the transcriptional regulators, hormone signaling, and cell proliferation/expansion.

In potato, the *miR166* family is amongst the most abundantly expressed miRNAs, with a total transcript per million exceeding 600,000, and exhibits tissue-specific and stolon-to-tuber stage-specific expression profiles in potato (Lakhotia *et al*., 2014; Kondhare *et al*., 2018). Further analysis of the small RNA sequencing data from Kondhare *et al*. (2018) revealed *miR166* to be differentially accumulated in stolons under long-day (LD; tuber non-inductive) and short-day (SD; tuber inductive) photoperiod. Here, we show that *miR166*-REVOLUTA-auxin module is an important regulator of tuber shape in potato, and modulates key agronomic traits including size, color, and productivity. *miR166* targets *StREVOLUTA*, a HD-ZIP class III transcription factor. StREV binds to the promoter of *StYUCCA7*, an auxin biosynthesis gene and activates its expression. Our work reveals a *miR166*–StREVOLUTA–StYUCCA7 function as one of the key regulatory pathways, providing mechanistic insights into the molecular control of tuber morphology and productivity.

## Results

### Identification, validation and expression analysis of *miR166* in Potato

Four precursors of *microRNA166* from the miRBase (the microRNA database; http://mirbase.org/) are predicted to form a secondary hairpin loop structure with mature *miR166* in its stem region (MFold; Zuker, 2003) (Figure S1A). To validate the presence of *miR166* precursors (precursors a, b, c and d) in potato, RT-PCR was performed and the amplified products were sequence confirmed (Figure 1A). Precursor-b was the least abundant form, as compared to the other three precursors.

**Figure 1.**
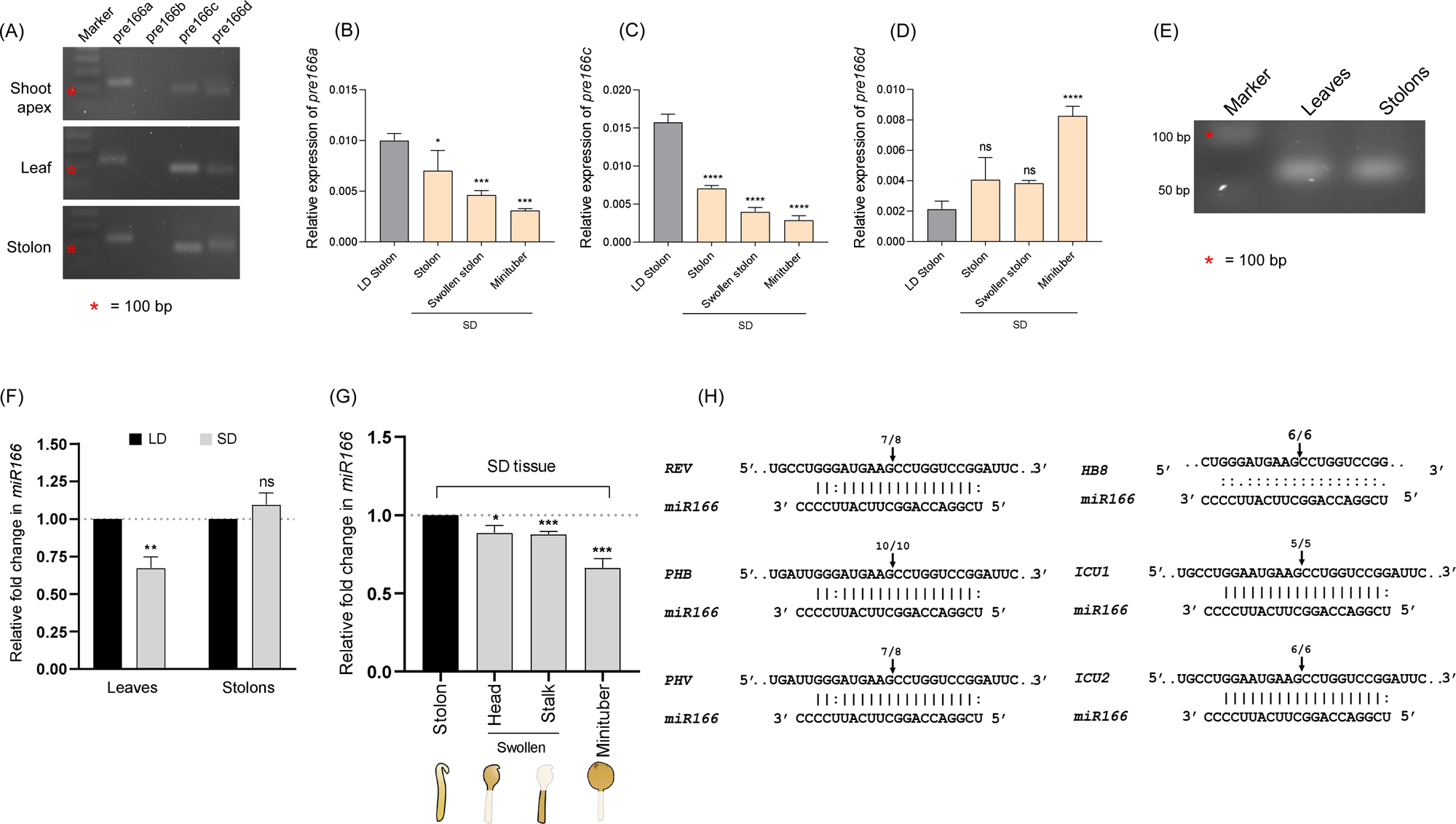
Identification, Validation, and Expression Analysis of *miR166* in Potato (A) RT-PCR of *miR166* precursors from shoot apex, leaves and stolons tissues of SD grown *Solanum tuberosum* ssp. *andigena*. (B-D) Quantification of precursor-a, -b, and -c levels from stolon-to-tuber transition stages (stolon, swollen stolon and mini-tuber) harvested from potato plants post 10 days of SD or LD induction. Relative transcript abundance of the gene expression in SD tissue is compared to LD stolons. Asterisks (one, two, three, and four) indicate significant differences at p < 0.05, p < 0.01, p < 0.001, and p < 0.0001, respectively, using Student’s t-test. ns = not significant at p > 0.05. *StEIF3e* was used as a reference gene. Detection (E) and relative fold change (F) of mature *miR166* from leaves and stolon tissues of SD grown (7 days post induction, 7 dpi) *Solanum tuberosum* ssp. *andigena* by stem-loop RT-PCR. (G) Relative abundance of *miR166* in stolon-to-tuber transition stages. *StU6* was used for normalization. Error bars indicate (±) SEM of three biological replicates, each with three technical replicates. Asterisks (one, two, three, and four) indicate significant differences at p < 0.05, p < 0.01, p < 0.001, and p < 0.0001, respectively, using Student’s t-test. ns = not significant at p > 0.05. (H) Confirmation of HD-ZIP III targets of *miR166*. Partial mRNA sequence of *ICU1*, *ICU2*, *REV*, *PHB*, *PHV*, and *HB8* aligned with *miR166*, numbers denote the fraction of cloned cleavage products that terminate at different positions (arrows).

Compared to LD, the levels of precursors-a and -c decreased in the stolons under SD photoperiod, while precursor-d increased (Figure 1B-D). The mature forms of all the precursors are identical, except for pre-*miR166*b, wherein it differs by a single base (Figure S1B). In our analysis of small RNA sequencing data from a previous study, we found *miR166* to be the most abundant microRNA in early stolon-to-tuber transition stages and exhibited differential accumulation under short-day (SD; tuber inductive) photoperiod (Kondhare *et al*., 2018) (Figure S2A-B). We detected the 21-bp mature *miR166* by stem-loop endpoint PCR from leaves and stolons of WT potato plants (Figure 1E). The amplified product was verified by sequencing, thus confirming that mature *miR166* is expressed in potato.

To determine whether photoperiod affects the accumulation of *miR166*, plants were grown in soil under long-day (LD; tuber non-inductive) and SD conditions. Stem-loop quantitative reverse transcription (qRT) -PCR analysis revealed a reduced accumulation of *miR166* in leaves under SD conditions, while stolons had comparable levels of *miR166* under both the photoperiods (Figure 1F). Stolons differentiating into tubers exhibited 10-40 percent reduction in the later stages of stolon-to-tuber transition (Figure 1G). *In silico* analysis suggested that *miR166* in potato potentially targets six *HD-ZIP III* members, similar to *Arabidopsis*. Using modified 5’ RLM-RACE, we mapped the cleavage site on the HD-ZIP transcripts (Figure 1H), confirming the targets of *miR166*. Overall, our results validate the existence of *miR166*-*HD-ZIP III* module in potato, and *miR166* showed tissue-specific accumulation under photoperiodic conditions.

### Knockdown of *microRNA166* causes pleiotropic phenotypic changes

Using a target mimicry approach, we generated stable transgenic lines of *miR166* in *S. tuberosum andigena ssp.* (Figure S3A). Of them, 2 independent lines (referred as KD #2 and KD #3) exhibiting 50-75% suppression of mature *miR166* were selected for further characterization (Figure 2A-B). *miR166* KD lines (MIM166) had reduced shoot height but number of nodes remained unchanged, implying a decreased inter-nodal distance, compared to the WT plants (Figure 2C-E). The leaf lamina from *miR166* KD lines appeared flat, hence we calculated the longitudinal and transverse curvature indices (CI^L^ and CI^T^) (Liu *et al*., 2010) to quantify the curvature. A curvature index of zero suggests that the leaf surface is completely flat. Both curvature indices, CI^L^ and CI^T^ of the KD lines were significantly reduced compared to WT (Figure 2F-G). The total chlorophyll content in the *miR166* KD lines were comparable to that of WT leaves; and leaves of KD #2 showed a slight increased anthocyanin accumulation, relative to the WT (Figure S3B). The vascular structures were distorted and had fewer xylem elements compared to the WT plants (Figure 2H). Further, an i*n vitro* root growth assay revealed that the MIM166 lines had decreased root length, shoot and root biomass, number of adventitious and lateral roots (Figure S3C-H). Thus, several phenotypic characters were altered upon constitutive and ubiquitous suppression of *miR166* in potato.

**Figure 2.**
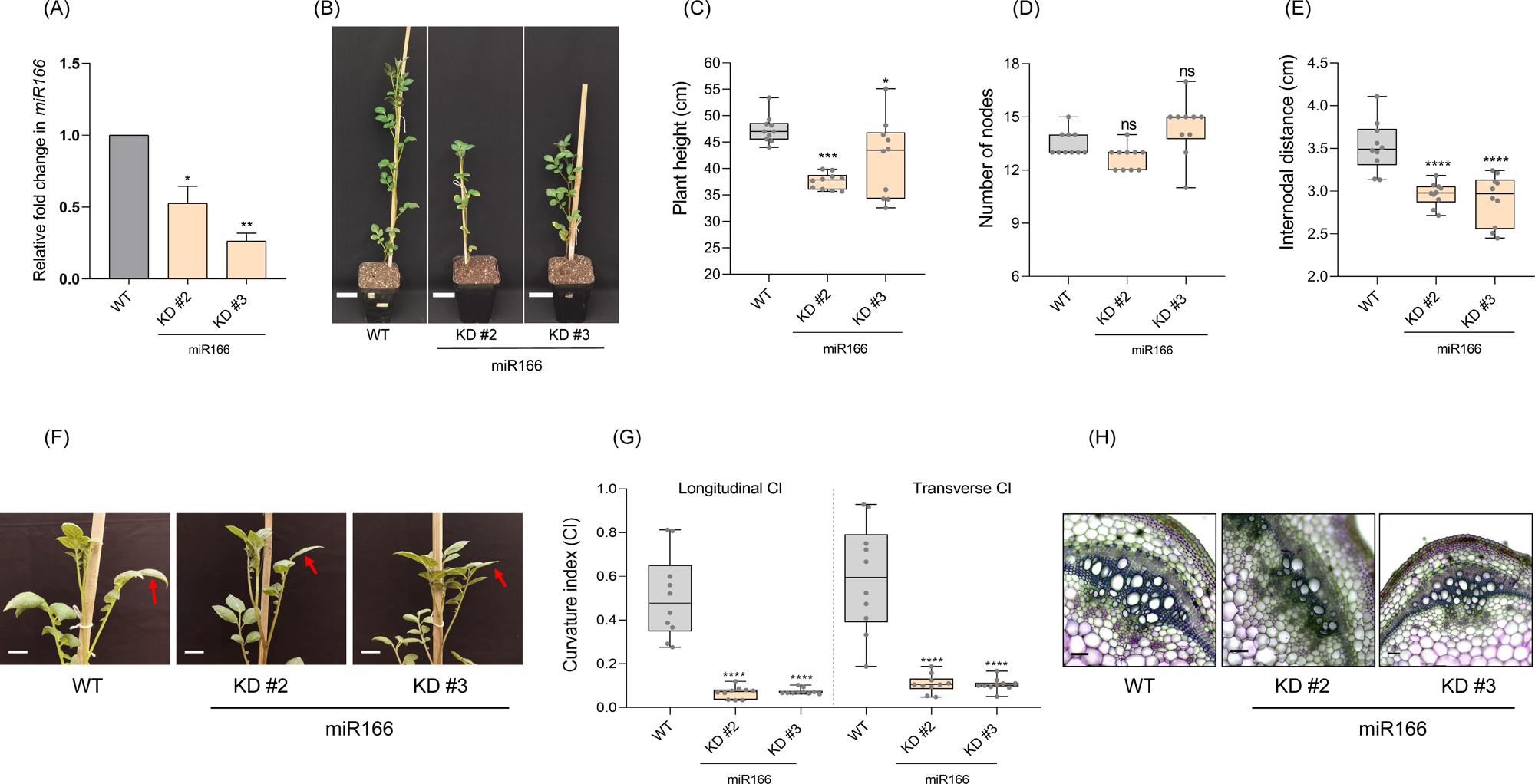
Knockdown of microRNA166 causes pleiotropic phenotypic changes. (A) Expression levels of *miR166* in mimicry lines (KD #2, KD #3) compared to WT. *StU6* was used as a reference gene. Error bars indicate standard error of means (SEM) from three biological replicates (n=3). Asterisks (one, two, three, and four) indicate significant differences at p < 0.05, p < 0.01, p < 0.001, and p < 0.0001, respectively, using Student’s t-test. ns = not significant at p > 0.05. (B) Plant architecture of *miR166* KD lines along with WT. Scale bar= 5 cm. Plant height (C), number of nodes (D) and internodal distance (E) of *miR166* KD lines in comparison with WT (n=10). (F) Photographs of 8-week old LD grown plants, showing reduced leaf curvature in KD plants (arrows). Scale bar= 2 cm. (G) The longitudinal and transverse curvature indices of leaves emerging from the 5th-node of 8-week old plants (n=10). Asterisks (one, two, three and four) indicate significant differences at p < 0.05, p<0.01, p<0.001 and p<0.0001, respectively, using one-way ANOVA. ns= not significant at <0.05. (H) Transverse sections of stem from *miR166* KD and WT LD grown plants. Scale bar= 50 microns.

### Suppression of *miR166* alters tuber shape, color and productivity

To examine the effect of *miR166* suppression on tuber development, we subjected the *miR166* KD lines to tuber inductive SD photoperiod in soil conditions. Compared to WT plants, the KD lines showed reduced tuber productivity in varying degrees, although the number of tubers per plant did not vary significantly (Figure 3A-B). Starch estimation assay showed no significant differences in the starch accumulation in tubers of any KD lines, compared to the WT tubers (Figure S3I). Interestingly, tubers from these KD lines were elongated as opposed to the WT spherical tubers (Figure 3C). The sphericity index for these tubers was greater than 1.5, while WT tubers had the sphericity index of around 1.0 (Figure 3D). However, the tuber volume of KD #2 was slightly reduced when compared to WT (Figure 3E). The pigmentation was observed only in the peel of the KD #2 and KD #3 tubers, while the flesh color was comparable to WT (Figure 3C). Metabolic profiling of the tuber peels revealed that KD #2 had increased accumulation of cyanidin 3-glucoside, while KD #3 had increased accumulation of both cyanidin 3-glucoside and pelargonidin 3-glucoside, relative to WT (Figure 3F). In contrast, delphinidin 3-glucoside was reduced in both the KD lines (Figure 3F).

**Figure 3.**
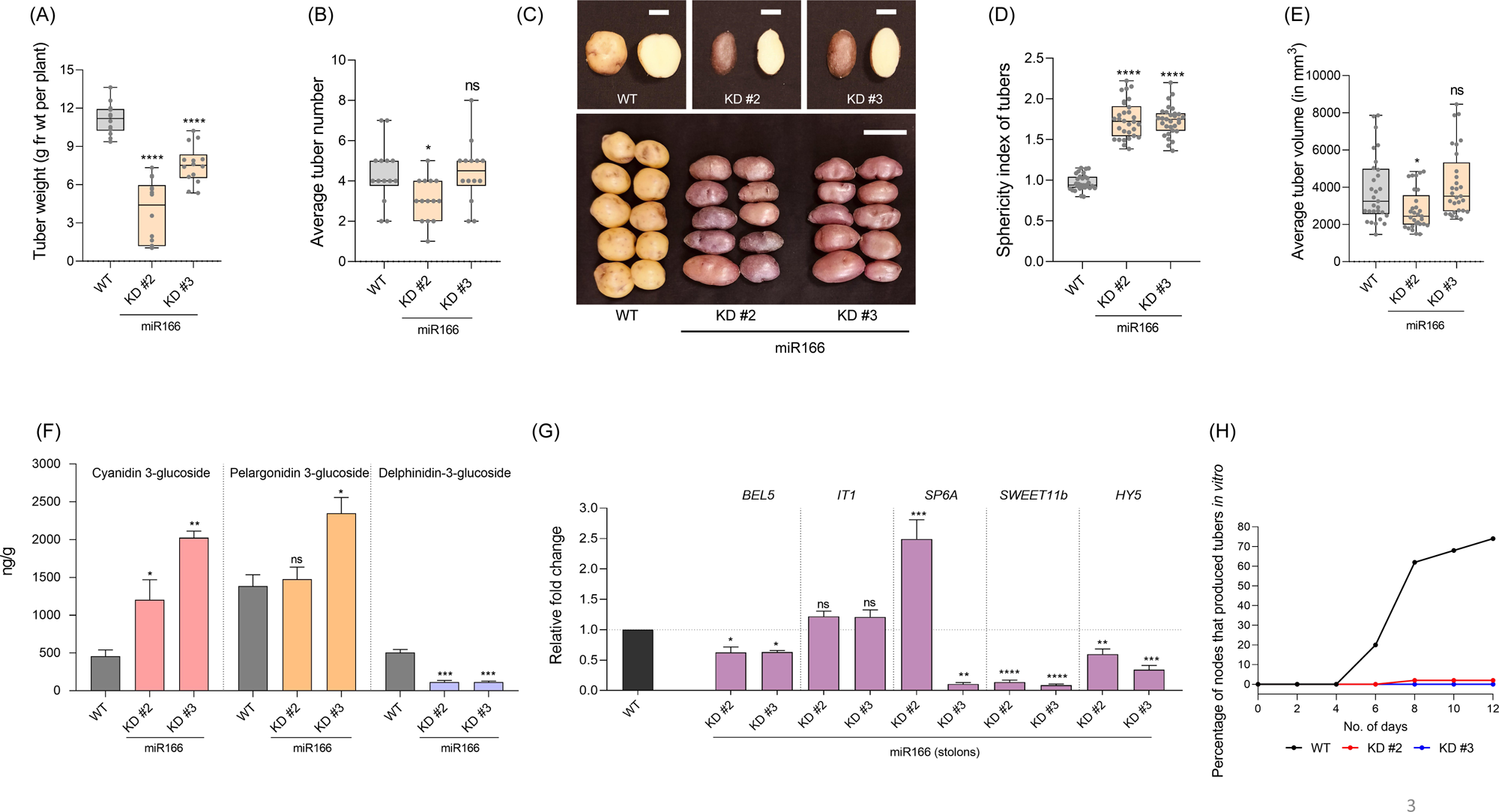
*miR166* knockdown lines have elongated, pigmented tubers with reduced tuber productivity (A) Tuber weight per plant and (B) average tuber number of *miR166* KD lines compared to WT. Data was taken from soil grown plants after two months of SD induction, and plotted from 14 individual plants per line. (C) Representative tuber images from *miR166* KD lines compared to WT, *miR166* KD tubers are elongated, and accumulate anthocyanin pigments in their peels. Cut tubers scale bar= 1 cm; Intact tubers scale bar= 3cm. The sphericity index (D) and average tuber volume (E) of *miR166* KD tubers in comparison to WT. The data is plotted for 30 tubers from each line. Asterisks (one, two, three and four) indicate significant differences at p < 0.05, p<0.01, p<0.001 and p<0.0001, respectively, using one-way ANOVA. ns= not significant at <0.05. (F) Quantification of cyanidin 3-glucoside, pelargonidin 3-glucoside and delphinidin 3-glucoside from the tuber peels of WT and *miR166* KD. The mean value (n = 4) ± SEM is presented. Asterisks (one, two, three, and four) indicate significant differences at p < 0.05, p < 0.01, p < 0.001, and p < 0.0001, respectively, using one-way ANOVA. ns = not significant at p > 0.05. (G) Relative fold change of key tuber marker genes in the stolons (7 days SD) of *miR166* KD, compared to the WT. Error bars indicate standard error of means (SEM) from three biological replicates (n=3). Asterisks (one, two, three, and four) indicate significant differences at p < 0.05, p < 0.01, p < 0.001, and p < 0.0001, respectively, using Student’s t-test. ns = not significant at p > 0.05. (H) *In vitro* tuber induction assay for *miR166* KD lines along with WT.

*BEL5* and *Identity of Tuber 1* (also known as *BRC1b*) are key positive regulators of tuberization (Chen *et al*., 2003; Tang *et al*., 2022). Relative quantification analyses from stolons of *miR166* KD lines showed a slight reduction in *BEL5* and no significant changes in *IT1* levels (Figure 3G). In potato, SP6A and SWEET11b are crucial for instrumenting symplastic sucrose loading under tuber-inductive conditions (Abelenda *et al*., 2019). *SWEET11b* was found to be significantly reduced in the stolons of both KD lines, along with varying levels of *SP6A* (Figure 3G). HY5 binds to the promoters to regulate the expression of *SWEET11* and *SWEET12* in *Arabidopsis* (Chen *et al*., 2016; Sakuraba and Yanagisawa, 2018). Stolons from KD plants had reduced *HY5* levels compared to the WT (Figure 3G). Further, we found that the tuberization potential of these KD lines was greatly reduced (80-100% reduction) under *in vitro* tuber inductive conditions (Figure 3H). Overall, the KD lines exhibited reduced tuberization potential *in vivo* and *in vitro*, accompanied with down regulation of key marker genes.

### RNA-sequencing analysis revealed differential expression of phytohormone-associated genes and accumulation of hormones in swollen *vs*. SSR region of tuberizing stolons

The phenotype of elongated and pigmented tuber was evident from the early stages of tuber development (Figure 4A). To identify the genes associated with the tuber phenotype, we carried out paired-end RNA sequencing on two distinct regions-the swollen region (Head) and the sub-swelling region (SSR)-from both WT and KD #3 samples (Figure 4B; Figure S4A). The WT_Head and SSR had 37 and 376 uniquely expressed genes respectively, and KD_Head and SSR had 115 and 120 uniquely expressed genes respectively (Figure 4C). A comparison between the Head *vs* SSR tissue revealed 8680 DEGs (4504 up- and 4176-down regulated) in the WT; while 7846 DEGs (3830 up- and 4016-down regulated) in the KD samples (Figure S4B-C; Table S1-S2). Using a cut-off of log2 fold change greater than 1 and p-value less than 0.05, KD_Head and SSR had 9774 and 10,209 DEGs compared to respective WT_Head and SSR tissues (Figure S4D-E; Table S3-S4).

**Figure 4.**
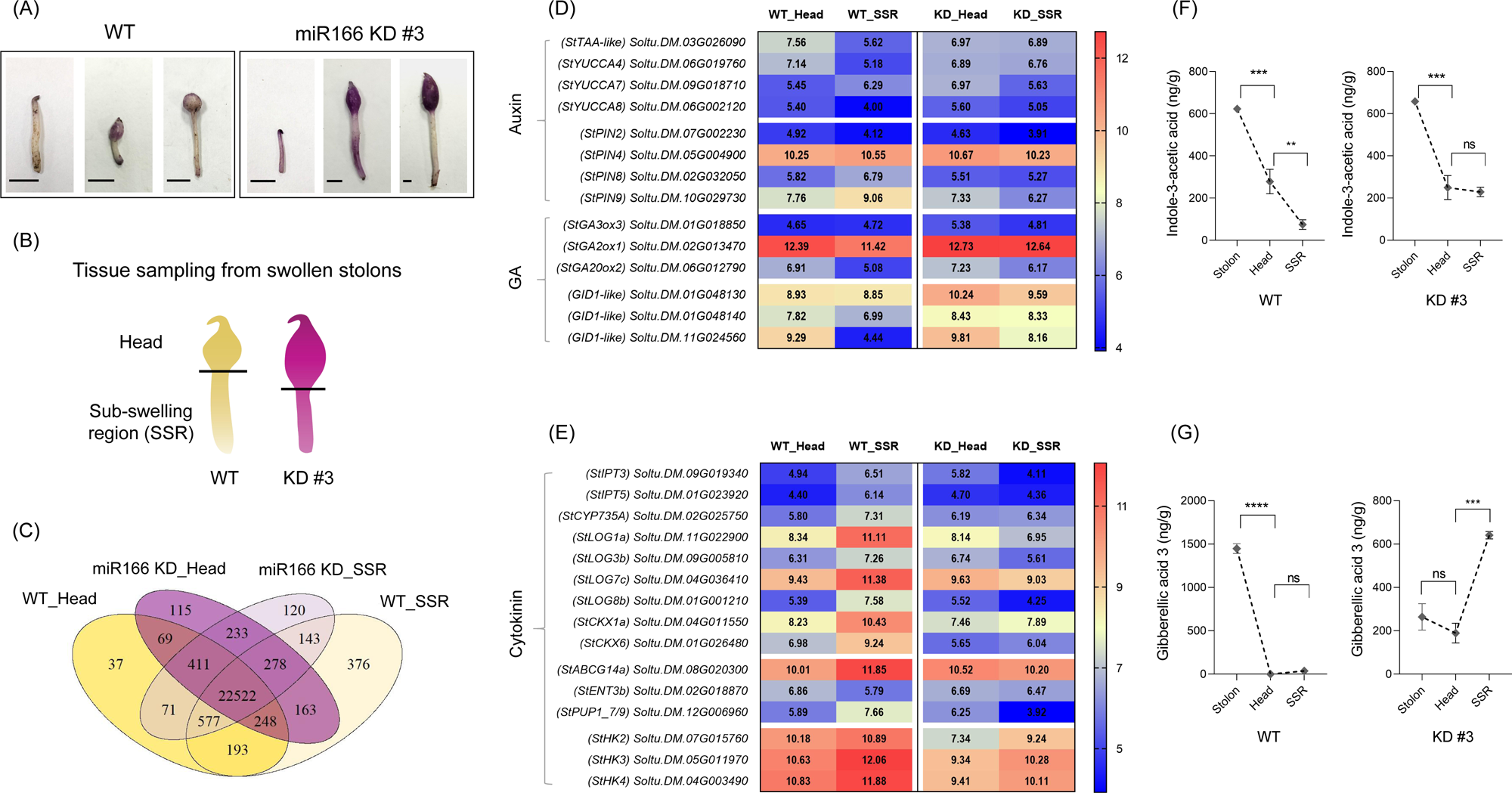
RNA-Seq analysis reveals differentially expressed genes in swollen and sub-swollen part of tuberizing stolons. (A) Stolon-to-tuber transition stages as seen in the WT and *miR166* KD #3. (B) A schematic illustration to depict the tissue collection carried out for RNA-sequencing analysis. The Head and sub-swelling (SSR) region were collected individually from WT and KD #3 stolons from plants grown under SD photoperiod for 12 days. (C) Venn diagram shows the summary of unique and shared genes expressed in KD #3 and WT tissues (Head *vs*. SSR). (D-E) The heat map represents differential expression of genes involved in Auxin, GA and CK metabolism and transport. Numbers inside the box represent variance stabilized transformed (*vs*T) values. Relative quantification of IAA-3 (F) and GA_3_ (G) from the Stolon, Head and SSR tissues from WT and *miR166* KD #3.

Hormone-related genes showed significant changes in their expression between the Head and SSR tissues from WT and KD (Figure 4D-E; Figure S5A-B). In WT, auxin biosynthesis genes - *StTAA-like*, *StYUCCA4* and *StYUCCA8* were up regulated in the Head compared to the SSR. This spatial expression distinction was lost in the KD line. Also, *StYUCCA5a* and *StYUCCA7* exhibited reversed expression patterns in KD line relative to WT. Auxin transport genes-*StPIN2/4/8/9* displayed markedly altered expression patterns in KD tissues, suggesting perturbed auxin distribution (Figure 4D; Figure S5A). The gibberellin (GA) biosynthesis genes *StGA3ox3* and *StGA20ox2* were found to be up regulated in both the KD tissues (Figure 4D; Figure S5A). Also, the differential expression of the GA catabolic gene *StGA2ox1*, which was evident between Head and SSR in WT, was abolished in the KD line. Expression of the cytokinin (CK) biosynthesis genes *StIPT3/5* and *StCYP735A* was altered and that of *StLOG1a/3b/7c/8b* and *CKX6* in the Head *vs.* SSR tissue was reversed in the KD lines (Figure 4E; Figure S5B). Cytokinin transporter *PUP1-7/9* showed significant reversed expression in the Head *vs.* SSR tissues of respective lines. RT-qPCR analysis of selected phytohormone-related genes validated the RNA-seq results, confirming the altered expression patterns (Figure S5C-D). In line with the transcriptomic data, auxin quantification using LC-MS/MS revealed similar auxin levels in both the Head and SSR regions of KD, whereas WT exhibited significantly lower auxin levels in the SSR (Figure 4F; Data S1.). Additionally, GA₃ levels in KD stolons were relatively 5.5 folds lower than those in WT, indicating an altered hormone dynamic in the KD line (Figure 4G).

In line with the reduced tuber productivity in MIM lines, positive regulators of tuberization such as *StBEL5* and *StCDF1* were slightly down regulated in KD, relative to WT. *StSWEET11b* levels were greatly reduced in both the KD tissues; while *StSP6A* and *IT1* were specifically down regulated in the SSR region (Figure S6A). RT-qPCR analysis of few marker genes such as *StBEL5*, *StSP6A* and *StHY5* corroborated with the RNA-seq data (Figure S6B-D).

### *miR166* overexpression exhibits minimal effects on vasculature and tuberization

*MicroRNA166* levels are abundant in the early stolon-to-tuber transitions and decrease at the mini-tuber stage (Figure 1G). To investigate the effects of *miR166* overexpression in potato, we generated stable transgenic lines overexpressing precursor *miR166-d* and evaluated their overall phenotype and tuberization potential under SD conditions. The overexpression (OE) lines OE #9 and #13 lines showed variable levels of mature *miR166* in leaf (Figure S7A-B). OE #13 exhibited a slight reduction in plant height and internodal distance, while the leaf curvature and overall architecture was comparable to those of WT plants (Figure S7C-F). The number of vascular cells in the OE #9 shoot seemed to be increased, relative to the WT plants (Figure S7G). Under SD conditions, the OE #9 exhibited a slight increase in tuber productivity, although the number of tubers remained largely unchanged (Figure S7H-J). RT-qPCR analysis revealed no significant difference in the levels of mature *miR166* in the stolons of OE lines, which may be due to the inherently high *miR166* in the WT stolon (Figure S7K). The tuber marker gene *BEL5* did not show significant variation between OE and WT stolons, while the tuber identity gene *IT1* was up regulated (Figure S7L). Additionally, auxin biosynthesis and transport-related genes were generally up regulated in the stolons of OE lines compared to WT (Figure S7M). Thus, *miR166* OE lines exhibited some phenotypic alterations, comparatively less severe than *miR166* KD lines.

### REVOLUTA, a *miR166* target affects tuberization

Using modified 5’RLM-RACE, we detected the *miR166* cleaved fragments of *StREV* as predicted by psRNAtarget finder (https://www.zhaolab.org/psRNATarget/home) and mapped the *miR166* cleavage site (Figure 1H; Figure 5A). In the WT stolons, REV transcripts were higher under long-day (LD) conditions and in shoot-forming stolons, whereas SD stolons exhibited dynamic expression across different tissues including the stolon, swollen region, SSR, and mini-tuber (Figure 5B). Stolons from *miR166* KD lines had increased *StREV* levels, while a slight reduction was observed in the OE stolons (Figure 5C). GUS assay of the stable transgenic lines expressing promREV::*uidA* showed promoter activity in the apical and axillary meristems, subtending regions likely corresponding to the vascular tissue, in the swollen regions of tuberizing stolons and at the apex of the mini-tuber stage (Figure 5D). The transcripts of *StREV* were localized in the adaxial side of developing leaves in the shoot apex of WT (Figure 5E). Stable transgenic lines of *StREV* anti-sense (*REV-AS*) exhibited variable degrees of *StREV* suppression in leaves and stolons (Figure S8A-B), however their overall growth and architecture was comparable to WT plants (Figure S8C). The stem vasculature in *StREV* AS #6 was smaller than that of in the WT (Figure S8D). Under SD photoperiod, all three AS lines displayed significantly reduced tuber yield, although the average number of tubers per plant remained largely unchanged (Figure 5F-H). Thus, *REVOLUTA* is a spatio-temporally regulated target of *miR166* in potato, and its suppression led to reduced tuber productivity.

**Figure 5.**
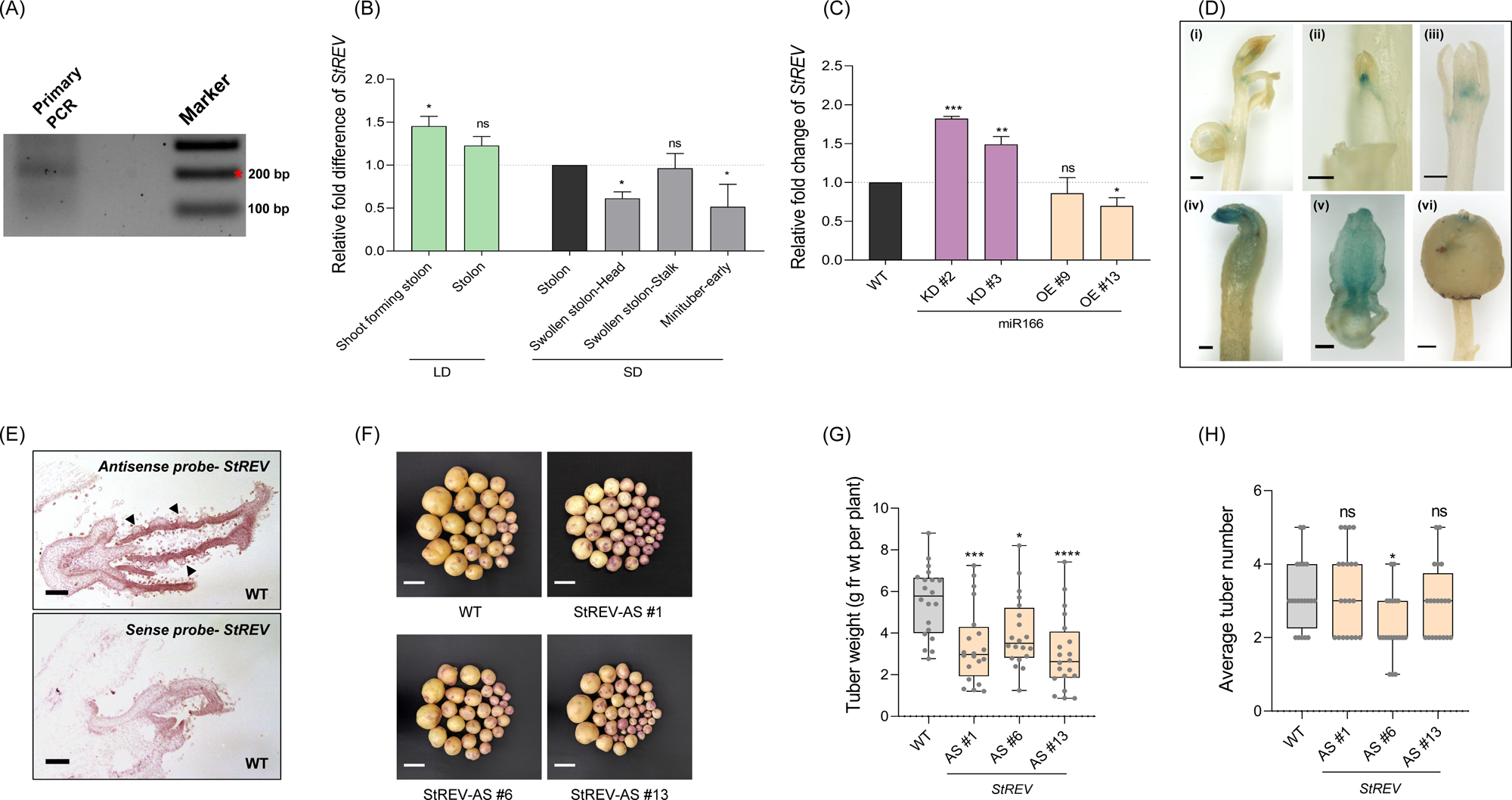
REVOLUTA affects tuberization. (A) Primary PCR product from modified 5′ RLM-RACE, to confirm *REV* as a target of *miR166* in potato. (B) Relative fold change of *REV* in differentiating stolons from WT plants. (C) Relative fold change of *REV* in stolons from *miR166* KD and OE lines. *StEIF3e* was used as reference gene. Error bars indicate standard error of means (SEM) from three biological replicates (n=3). Asterisks (one, two, three, and four) indicate significant differences at p < 0.05, p < 0.01, p < 0.001, and p < 0.0001, respectively, using Student’s t-test. ns = not significant at p > 0.05. (D) Promoter expression of *StREV* in *promStREV:uidA* transgenic lines. (i) *in vitro* plant, (ii) axillary node, (iii) shoot-forming stolon, (iv) swollen stolon, (v) T.S. of swollen stolon, and (vi) mini-tuber. Scale bar= 1000 um for (i) and (vi); Scale bar= 500um for (ii)-(v). (E) Longitudinal sections of shoot apex from WT plants, black arrow heads show polarized localization of REV transcripts in developing leaves. Scale bar= 100 um. Tuber productivity and tuber numbers (F-H) from REV AS lines, relative to the WT. Scale bar= 2 cm. Asterisks (one, two, three and four) indicate significant differences at p < 0.05, p<0.01, p<0.001 and p<0.0001, respectively, using one-way ANOVA. ns= not significant at <0.05.

### StREVOLUTA activates *StYUCCA7*

The AT[G/C]AT palindromic sequence was identified as the *in vitro* binding sequence for HD-ZIP III proteins, through which they bind the promoters of auxin-related genes in *Arabidopsis* and regulate their expression (Sessa *et al*., 1998). *REVOLUTA* is known to up regulate the promoter activity of auxin-biosynthesis genes TAA1 and YUCCA5 in Arabidopsis (Brandt *et al*., 2012). Our *in-silico* promoter analysis of auxin biosynthesis genes in potato revealed the presence of these core binding motifs within their upstream regulatory regions, with maximum motifs being present on the promoter of *StYUCCA7* (Figure 6A). RT-qPCR from the stolons of stable transgenic lines of *StREV* suppression showed reduced accumulation of *StYUCCA* transcripts (Figure 6B), suggesting that the biosynthesis genes could be the targets of StREV. To test this hypothesis, we performed dual-luciferase assay in *Nicotiana benthamiana* leaves. When promoter *StYUC7* was co-infiltrated with *5’m-StREV-FLAG*, the Firefly/Renilla ratios were 10-fold higher that when infiltrated alone, or in presence of an empty vector, suggesting that StREV up-regulated *StYUC7* at the transcriptional level (Figure 6C). To determine whether StREV binds directly to the promoter of *StYUCCA7*, yeast one-hybrid assay was performed. StREV could activate the promoter of *StYUC7* as seen by the robust growth of mated colony on increasing concentrations of 3-AT till 80mM, after which the growth was arrested on higher concentrations (Figure 6D). Thus, StREV could bind to the promoter and up regulate the expression of the auxin biosynthesis gene *StYUCCA7*.

**Figure 6.**
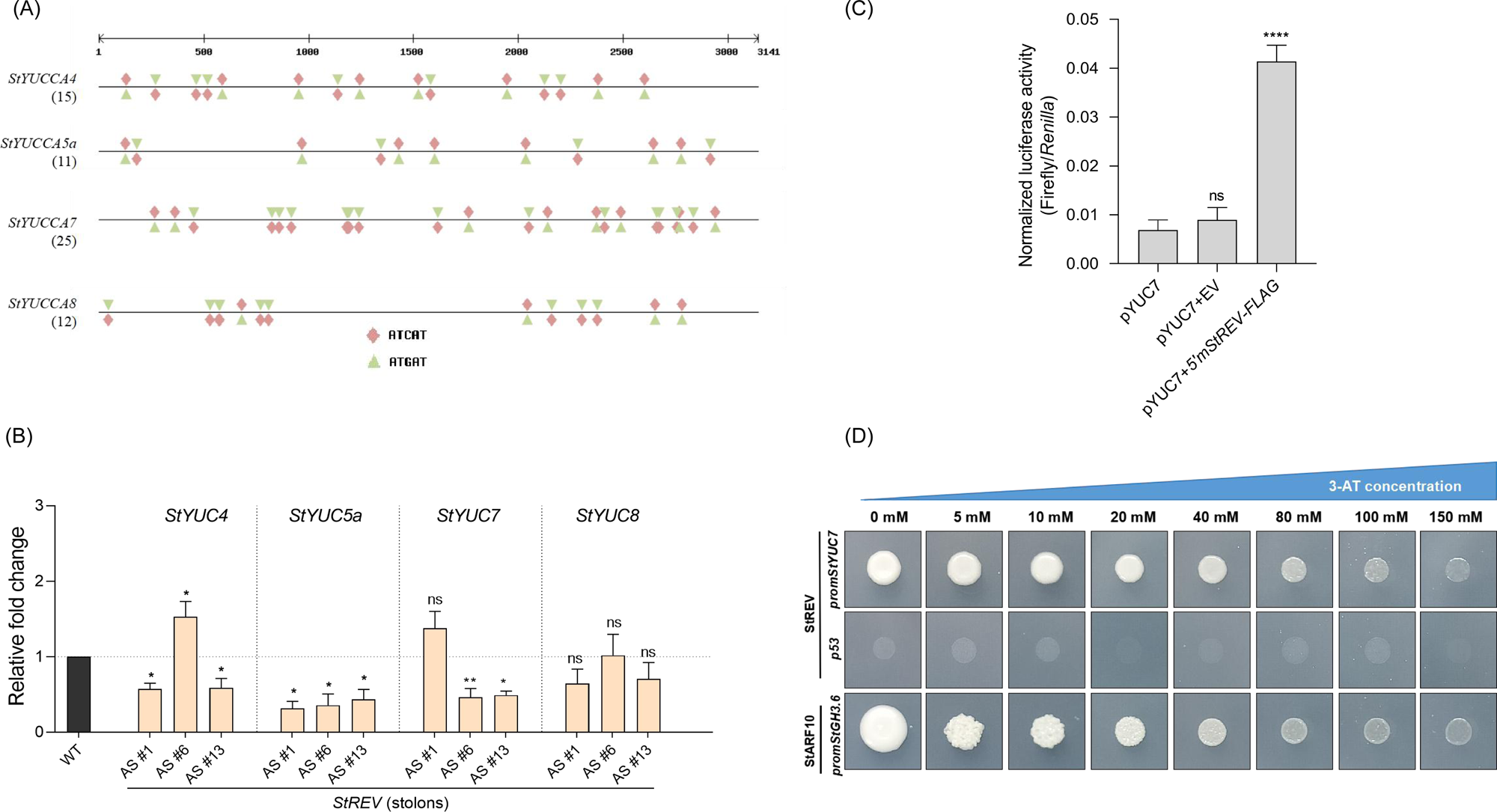
(A) Promoters of auxin biosynthesis genes harbor the AT(C/G)AT sequence, a core motif of HD-ZIP III binding site. (B) RT-qPCR of *REV*-AS stolons show reduced transcript abundance of auxin biosynthesis genes. Dual luciferase assay (C) shows approximately 10 fold upregulation of promoter *StYUC7* in presence of StREVOLUTA. A *5’m-StREV-FLAG* construct was used that is resistant to cleavage by *miR166*. Asterisks (one, two, three and four) indicate significant differences at p < 0.05, p<0.01, p<0.001 and p<0.0001, respectively, using one-way ANOVA. ns= not significant at <0.05. (D) Yeast-mated clones containing *StREV* prey protein and prStYUC7 bait, or StARF10 prey protein and *prStGH3.6* bait (positive control) grew in media containing up to 80 mM 3-amino-1,2,4-triazole (3-AT), indicating the binding of StREV to *StYUC7* promoter. The p53 binding site was used as a negative control. The yeast-mated clones harboring StREV and *p53* did not grow on the selection media, suggesting that there is no interaction between the *p53* binding site and StREV protein.

## Discussion

Potato is a well-studied model for molecular and hormonal control of tuberization, however, limited knowledge is available on the molecular mechanisms that regulate tuber morphology. In this work, we investigated the role of *miR166* in tuber development in *Solanum tuberosum* spp. *Andigena* 7540. *MiR166* targets six members of the HD-ZIP class III transcription factor family, and is highly expressed in differentiating stolons under SD photoperiod. Suppression of *miR166* led to the pleiotropic effects on overall plant development, particularly affecting tuber shape, color and productivity. Further, StREVOLUTA, a HD-ZIP class III TF regulates auxin biosynthesis during early stolon-to-tuber differentiation *via StYUCCA7.* Our findings identify *miR166*-HD-ZIP III as an important regulator of tuber shape, color and productivity and provides molecular insights about morphology of the storage organ.

### The functional roles of *miR166* are conserved in potato

MicroRNAs are key regulators of plant development and defense responses, and amongst them, *miR166* represents a well-conserved family that targets members of the HD-ZIP III transcription factors. Despite the diverse roles attributed to *miR166* in several plants’ development, productivity, and stress responses (Ding *et al*., 2013; Zhang *et al*., 2024), its functional characterization in potato and relevance in storage organ development remains unexplored. In our preliminary analysis of small RNA sequencing datasets (Lakhotia *et al*., 2014; Kondhare *et al*., 2018), we identified *miR166* as the most abundant microRNA in potato stolons, suggesting its involvement in stolon-to-tuber transition stages. A constitutive and ubiquitous knockdown of *miR166* in potato (MIM166) led to the development of several pleiotropic phenotypes. The MIM166 lines had reduced plant height, having no significant changes in the number of nodes (Figure 2C-E). In soybean, Zhao *et al*. (2022) showed that suppression of *miR166* led to decreased plant height. ATHB14-LIKE (a HD-ZIP III TF) promoted the expression of a GA catabolic gene-*GIBBERLLIN 2 OXIDASE2*, causing decreased bioactive GA levels in the plant affecting the height. In potato, MIM166 *in vitro* shoots and roots had increased levels of inactive GA8 (degraded product) and elevated expression of *GA2ox1* in stolons, compared to the WT tissues (Figure S9A-B), suggesting a similar GA dynamics in potato. In *Arabidopsis,* miR165/166 suppression led to pleiotropic effects on leaf development such as trumpet-shaped leaves, ectopic leaflets on the abaxial side, increased leaf numbers and accumulation of purple pigment (Jia *et al*., 2015). Knockdown lines of *miR166* in rice exhibited a rolled-leaf phenotype, having smaller bulliform cells and abnormal sclerenchymatous cells (Zhang *et al*., 2018). In potato, MIM166 lines exhibited reduced curvature of the leaf blades, variations in the total leaflet length and number of lateral leaves (Figure 2F-G; Figure S9C). Thus, *miR166* suppression causes diverse leaf phenotypes in different plants, suggesting a conserved yet species-specific role in regulating leaf development. Two targets of *miR166*, ATHB15 and HB8, play antagonistic roles in the regulation of xylem development in *Arabidopsis* (Baima *et al*., 2001; Kim *et al*., 2005). *ATHB15* is exclusively expressed in vascular tissues and its suppression leads to expanded xylem tissue and inter-fascicular region, indicative of accelerated vascular cell differentiation from cambial/procambial cells (Kim *et al*., 2005). We observed that stable potato transgenic lines of *miR166* (MIM and OE) had altered vascular structures in the stem (Figure 2H; Figure S7G), suggesting *miR166*-*HD-ZIP III* network might also influence differentiation of vascular cells in potato. Singh *et al*. (2014) showed that phytohormones dynamically regulate the expression of miR165/166 in roots of *Arabidopsis*, and control root meristem size and growth. *In vitro* root growth assay of potato, MIM166 lines showed decreased root length, root biomass, number of adventitious and lateral roots (Figure S3C-H), suggesting the important role of *miR166* in root development of potato. Overall, these pleiotropic phenotypes of the potato MIM166 lines recapitulate the phenotypic traits observed in other plants, and highlight its conserved functional roles as a central regulator of growth and development.

### *miR166* impacts tuber shape, color, and productivity in potato

Phytohormone quantification, RNA sequencing of tuberizing stolons (Head *vs.* SSR), and anatomical and molecular analyses revealed molecular insights into the altered tuber phenotypes of MIM166 plants. In potato stolons, high GA levels inhibit tuberization, while auxin content increases under inductive conditions. In MIM166 stolons, the GA catabolic gene *GA2ox1* was up regulated (Figure 4D), accompanied by a 5.5-fold reduction in bioactive GA₃ (Figure 4G). Genes involved in auxin biosynthesis (*StTAA-like, StYUCCA4/5a/7/8*) and transport (*StPIN2/4/8/9*) showed altered expression in KD_Head *vs*. WT_Head, suggesting disrupted auxin distribution. Consistently, auxin levels in KD Head *vs*. SSR were comparable, unlike the distinct accumulation in WT (Figure 4D). Cytokinin, which is essential for cell division and expansion during tuber bulking (Mingo-Castel *et al*., 1976; Sattelmacher and Marschner, 1978), had reduced expression of its biosynthesis and transport genes in SSR (Figure S5B), affecting the radial cell expansion. Moreover, MIM166 tuber sprouts were stunted, pigmented, and highly branched (Figure S10A), indicating an altered auxin–CK dynamics. We propose that higher auxin in the stalks of developing tubers, altered polar auxin transport, and disrupted auxin–GA–CK balance in the differentiating stolons caused anisotropic expansion leading to elongated tubers in MIM166 plants.

Cyanidin 3-glucoside, pelargonidin 3-glucoside, and delphinidin 3-glucoside are water-soluble anthocyanins with strong antioxidant and anti-inflammatory properties (Noda *et al*., 2002; Duarte *et al*., 2018). In our study, MIM166 tubers showed differential accumulation of these pigments in the peel, but not in their flesh (Figure 3C). A similar phenotype was reported in tomato, where overexpression of a petunia chalcone isomerase (CHI) gene increased peel flavonoids by 78-fold without affecting the flesh (Muir *et al*., 2001). In our transcriptomic analysis, we observed an up regulation of multiple structural and accessory genes from the anthocyanin biosynthesis pathway in KD tissues compared to WT, consistent with the increased pigment accumulation (Figure 3F; Figure S10B). Transcription factor families which highly correlated with these pathway genes-bHLH, WRKY, GRAS, bZIP, LBD, ARF, B3, G2-like, HD-ZIP, MYB, and MYB-related also showed significant differential expression (Table S3–S4; Figure S10C). In apple, auxin-induced degradation of MdIAA121 releases MdARF13, which functions as a negative regulator of the anthocyanin biosynthetic pathway (Wang *et al*., 2018). Similarly, auxin suppresses anthocyanin accumulation by promoting the degradation of MdIAA26 (Wang *et al*., 2020). In potato, tuberizing stolons have high auxin levels. We speculate that this high auxin could have restricted pigment accumulation to the tuber peel in the MIM166 lines, thereby resulting pigmented peels but non-pigmented tuber flesh. Conversely, flavonoids have been reported to act as inhibitors of auxin transport (Gayomba *et al*., 2016), which may have altered auxin distribution in the KD stolons. Further studies are required to ascertain how auxin dynamics in tuberizing stolons influence anthocyanin accumulation and, consequently, tuber morphology.

Under tuber inductive SD photoperiod, SWEET11b-SP6A interacts to mediate the symplastic loading of sucrose, and prevents its loss to the apoplastic route (Abelenda *et al*., 2019). MIM166 plants showed strong reduction in SWEET11b levels (Figure 3G; Figure S6A), and had distorted vascular structures (Figure 2H), both of which could have contributed to impaired sucrose transport causing reduced tuber productivity, while not significantly affecting the starch content per gram fresh weight of the tuber (Figure 3A; Figure S3I). Interestingly, MIM166 plants developed floral buds at the shoot apex and aerial stolon/tubers under SD photoperiod (Figure S11A-C), denoting the formation of ectopic sinks due to the improper sucrose distribution and/or cytokinin metabolism. As SD induction progressed, the belowground developing tubers acted as the primary and stronger sinks, thus limiting sucrose availability for the ectopic sinks leading to floral bud abortion. Thus, the reduced belowground tuber productivity could be a cumulative effect of altered vascular structures, improper symplastic loading of sucrose, and impaired tuber-bulking at the stolon apex due to the disrupted phytohormone dynamics.

### *MiR166*-Revoluta-auxin: Role in tuberization

Auxin plays a key role in the stolon-to-tuber transition process. The stolon apical meristem (StAM) is a major site of auxin biosynthesis, where YUC-like1 (*StYUC4*) acts as a key contributor (Roumeliotis *et al*., 2012). In *Arabidopsis*, REVOLUTA regulates promoters of auxin biosynthesis, transport, and signaling genes (Brandt *et al*., 2012; Reinhart *et al*., 2013). In our study, phenotypes such as impaired root growth, increased tuber sprout branching, and altered auxin-related gene expression suggested a role for REVOLUTA in potato (Figure 5A–C; Figure S3C-H; Figure S10A). Through GUS assay, we observed that the promoter activity of *StREV* was visible in the stolon apex, vascular region and subsequent tuberizing stages (Figure 5D). This was similar to the activity of an auxin responsive promoter *DR5:GUS* observed by Roumeliotis *et al*. (2012), suggesting a putative overlap of *StREV* expression and auxin accumulation. *StREV* antisense (REV-AS) lines exhibited reduced tuber yield (Figure 5F-H). Also, StREV could bind to the promoter of *StYUC7* and activate its expression (Figure 6D-E). These findings suggest a hitherto unknown role of REV in tuberization, probably through auxin.

We propose a model for the *miR166*–REV–auxin regulatory module in influencing tuber shape and size in potato (Figure 7). In WT, high levels of *miR166* in stolons fine tune the expression of *StREV*. This leads to a balanced metabolism of auxin in the tuberizing stolons, resulting in isotropic expansion and formation of spherical tubers. In the MIM166 lines, suppression of *miR166* caused increased and/or ectopic expression of *StREV*. This altered the auxin biosynthesis, and polar auxin transport, causing anisotropic expansion of the stolons and forming elongated tubers. In the *REV-AS* lines, lower levels of *StREV* affected auxin biosynthesis, and resulted in reduced tuber size and lower productivity compared to the WT. Collectively, these findings support a model, wherein the *miR166*–REV–auxin regulatory module governs auxin biosynthesis during stolon differentiation, thereby influencing tuber morphology (Figure 7). Given that auxin levels are highly dynamic during tuberization, further studies are needed to elucidate the spatio-temporal regulation of the *miR166*–REV and its impact on auxin synthesis/transport. Additionally, members of the HD-ZIP III TF family could also play a role in the stolon-to-tuber transitions and needs further investigations (Figure S12A-B).

**Figure 7.**
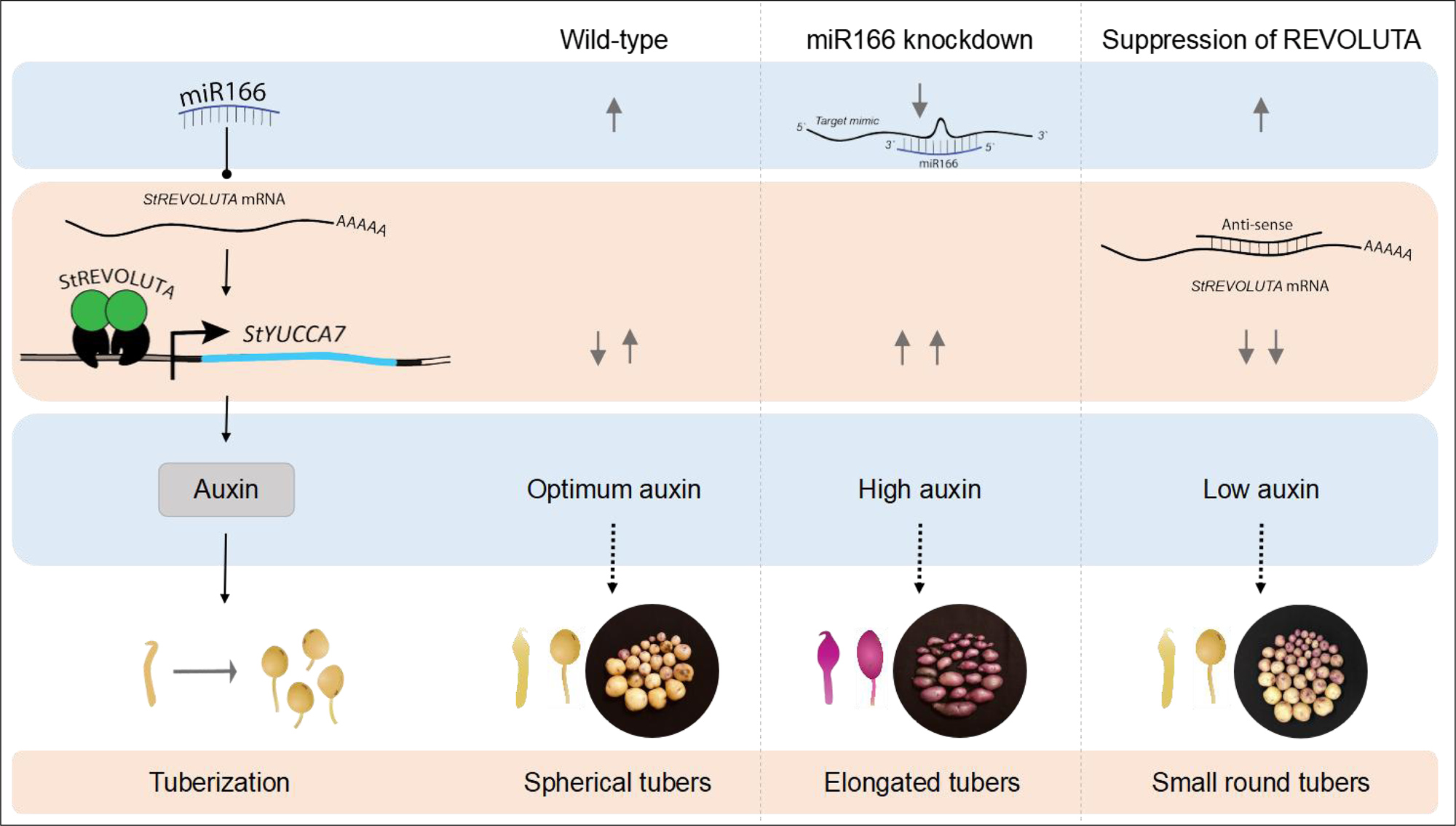
A proposed model for the role(s) of *miR166*-REVOLUTA-auxin module in potato tuberization. *miR166* in potato targets the *StREVOLUTA*. StREV activates the expression of auxin biosynthesis gene-*StYUCCA7*. In MIM166 line, decreased levels of *miR166* leads to increased/ectopic *REV* expression, causing perturbed auxin biosynthesis and transport leading to the development of elongated tubers. In *StREV*-antisense, the tubers are spherical, with reduced size and productivity.

Our transcriptomic datasets revealed that genes located within QTL regions on chromosomes 5, 10, and 12-associated with tuber shape and eye depth-were differentially expressed (Figure S13A; Tables S3–S4). Though we do not see a marked difference in the *StOFP20* expression (Ju *et al*., 2023); differential expression of 19 other OFPs suggests the involvement of more than one OFP genes in influencing tuber shape (Figure S13B). Additionally, several genes within the *Ro* locus (Fan *et al*., 2022) were down regulated in KD_Head compared to WT_Head (Figure S13C; Table S3). Further studies of these genes will elucidate their role(s) in determining tuber morphology.

Tuber morphology is shaped by the coordinated interplay of phytohormone and developmental gene regulatory networks that govern cell division, expansion, and tissue organization. Although the *miR166*-HD-ZIP III module is widely associated with adaxial-abaxial polarity in dorsiventral lateral organs, our findings uncover its functional relevance in a radially symmetric storage organ. We demonstrate that this regulatory module contributes to tuber development by modulating auxin-associated pathways, with REVOLUTA regulating auxin biosynthesis genes and influencing tuberization. Further studies in functional diversification of HD-ZIP III members will provide insights into how small RNA-transcription factors can fine-tune organ architecture in diverse developmental contexts.

## Materials and Methods

### Plant material and growth conditions

A photoperiod-responsive potato cultivar *Solanum tuberosum* ssp. *andigena* 7540 which tuberizes under SD conditions, was used in this study. *In vitro* plants (wild-type and transgenic lines) were propagated from axillary node sub-cultures on Murashige and Skoog’s basal medium with 2% (w/v) sucrose (Murashige and Skoog., 1962) and were grown in a plant growth incubator (Percival Scientific, Inc.) with a light intensity of 300 μmol m−2 s−1 under LD conditions unless mentioned otherwise.

### Prediction and detection of *miR166* precursors in potato

Four precursor sequences (stu-*MIR166*a, stu-*MIR166*b, stu-*MIR166*c and stu-*MIR166*d) of *miR166* were retrieved from the miRBase (http://mirbase.org/; Griffiths-Jones S *et al*., 2008). The secondary structures of these precursors were predicted using the Mfold software (http://www.unafold.org/mfold/applications/rna-folding-form.php; Zuker, 2003), and structures with lowest free energies were then further screened for having the characteristic stem-loop secondary structure with the mature miRNA sequence in the arm region of the stem-loop. All four miRNA precursors were then validated from *S. tuberosum* ssp. *andigena* 7540 by RT-PCR analysis. The sequences of all the oligos used in this study are provided in Table S5, and the IDs for all the genes are in the Table S6.

Total RNA was harvested from shoot apex, mature leaves and stolon tissues of potato using RNAiso Plus (DSS TAKARA) as per the manufacturer’s instructions. cDNA was synthesized using two microgram (2µg) of total RNA, Superscript IV Reverse Transcriptase (Invitrogen), and oligo dT primers. miRNA precursors were amplified by RT-PCR using precursor-specific primer pairs. The amplicons were cloned into sub-cloning vector pGEMT-easy (Promega) and sequence verified.

### Validation of mature *miR166*

Total RNA was isolated using RNAiso Plus (DSS TAKARA) from mature leaves and stolons of WT potato plants. One microgram (1 µg) of total RNA was used for cDNA preparation using *miR166* specific stem-loop primers (STP) followed by their end-point PCRs using *miR166* specific forward and universal reverse primer as described in Varkonyi-Gasic *et al*. (2007) (Table S5).

### Tissue-specific relative abundance of *miR166* under SD and LD conditions

For investigating the influence of photoperiod on tissue-specific accumulation of mature *miR166*, *in vitro* grown wild-type *S. tuberosum* spp. *andigena* 7540 plants were transferred to soil and maintained under LD photoperiod with 300 μmol m−2 s−1 light intensity for a period of ten weeks (until they attained 10-12 leaf stages) in a growth chamber (Percival Scientific, Inc.). Later, half of the plants were transferred to tuber-inductive SD photoperiod for 7 days, while the remaining plants were maintained under LD conditions. Leaves and stages of stolon-to-tuber transitions were collected at 7 days post LD/SD induction in triplicates. Isolation of total RNA and cDNA synthesis was carried out as described before. qPCR reactions were performed on CFX96 Real-Time System (BIO-RAD) with *miR166*-specific forward primer and universal reverse primer, using TAKARA SYBR green master mix (Takara-Clontech). The reaction mix was incubated at 95 °C for 30 s, followed by 40 cycles at 95 °C for 5 s, gene-specific annealing temperature for 15 s and extension for 72 °C for 15 s. PCR specificity was checked by melting curve analysis, and data were analyzed using the 2^-ΔΔCt^ method (Livak and Schmittgen, 2001).

### Generation of constructs and potato transgenic lines

MicroRNA166 overexpression (OE) and mimicry (MIM) constructs for overexpression and suppression of the *miR166*, respectively, were previously generated in our lab. Briefly, *miR166* OE construct-35S:St-pre166D-pBI121 was generated by amplifying precursor (St-pre166-d) from potato RNA using primers Pre166d-FP and Pre166d-RP (Table S5). The amplified product (108 bp) was cloned into the binary vector pBI121 downstream to the 35S CaMV constitutive promoter. For knockdown (KD) construct, we used artificial target mimicry (MIM) strategy. The KD construct (originally in pGREEN vector) was obtained from European *Arabidopsis* Stock Centre (NASC) (Todesco *et al*., 2010), and the MIM166 insert (542 bp) was re-cloned to pBI121 binary vector to generate 35S:MIM166-pBI121. Both these constructs-OE and MIM were then mobilized into *Agrobacterium tumefaciens* strain GV3101. Transgenic potato plants in *S. tuberosum* spp. *andigena* 7540 were generated as described in Banerjee *et al*., 2006. Kanamycin-resistant transgenic plants were further screened by molecular methods and selective lines were used for further experiments.

After 4 weeks of *in vitro* growth, potato plants were transferred to soil, grown for 7-9 weeks under LD conditions, before initiating SD induction. Phenotypic characteristics such as plant height, number of nodes and leaf curvature were recorded just before the initiation of SD photoperiod treatment, whereas the frequency of floral bud formation and tuber productivity parameters were measured after 1 and 8-9 weeks of SD induction, respectively.

### Phenotypic analysis

All the phenotypic analyses were performed on 6–7 weeks old soil-grown plants. Shoot height was measured from the bottom-most node up to the plant apex. For measurement of leaf curvature indices, mature terminal leaves at the fifth node were analyzed as described in Liu *et al*., 2010. These experiments were performed at least three times with similar results.

### Assessment of tuber productivity

One-month-old plantlets of the transgenic lines and WT were transferred to soil in small round pots and hardened for one week under LD conditions in a growth chamber (Percival Scientific, Inc.) with a light intensity of 300 μmol m−2 s−1, day temperature of 24°C, and night temperature of 22°C. This was followed by an additional three weeks of incubation in the same pots. Subsequently, the plants were re-potted into medium-sized pots and continued to grow under LD conditions for an additional five weeks. When the plants reached the 10-12 leaf stage, they were transferred to tuber inductive short-day (SD) conditions (16h dark and 8h light) with day and night temperatures of 22°C and 20°C, respectively. A subset of these plants was harvested after 7 days of induction, and stolons were collected for RT-qPCR analysis. After 60 days of induction, remaining plants were evaluated for their tuber productivity (gram fresh weight [g fr wt] per plant) and tuber numbers. This experiment was carried out 3 times, with consistent results. For tuber sphericity index and volume assessment, 30 tubers per line were measured for their diameters (Diameter1, Diameter2 and height). *In vitro* tuber induction assay was performed (experiment was repeated three times independently) as described in Kondhare *et al*., 2021.

### Starch estimation

Starch estimation was carried out twice, independently, as described in Kamble *et al*. (2023), Hawkins *et al*. (2021) and Kringel *et al*. (2020), with minor modifications. Tuber tissue was well-homogenized in cold water containing 0.35% sodium metabisulphte. The slurry was filtered to remove the debris. Filtered solution was kept undisturbed for 1 hour at 4 degrees. Insoluble material was collected by centrifugation, followed by 3 washes with 80% ethanol, and then finally re-suspended in water. Starch was digested using alpha amylase (SRL), glucoamylase (SRL) and Amyloglucosidase (Sigma). Glucose was estimated using the DNSA method, and compared with a standard graph prepared using Starch (Sigma), digested under identical conditions.

### RNA sequencing

For RNA-sequencing, *miR166* KD #3 and WT plants were grown in soil for 10-weeks under LD conditions and subjected to SD induction for 12 days. Swollen stolons were harvested in triplicates (from at least 20 plants per replicate) and divided into swelling (Head) and sub-swelling region (SSR) and snap-frozen in liquid nitrogen. The tissues were crushed using liquid Nitrogen and stored in −80⁰ C until further use. For RNA sequencing and RT-qPCR purposes, 100 mg of tissue from each sample was used to isolate the total RNA using RNAiso Plus as per the manufacturer’s instructions. RNA quality and integrity were assessed, and RNA-sequencing was performed by Novelgene Technologies Pvt. Ltd., Hyderabad. Paired-end sequencing was performed on Illumina NovaSeq 6000, adapter sequences and low-quality reads were filtered using FASTP to get high-quality reads on the parameter of <Q20, length < 50. Further processing was performed as described in Kumar *et al*. 2020. Validation of selected target genes identified in the RNA-seq analysis was done using qPCR as described above.

### Detection of phytohormones and pigments by LC-MS/MS

Tissues-stolon, swollen stolon (Head and SSR), tuber peel, tuber flesh were harvested, crushed in liquid Nitrogen, and stored at −80⁰ C until further use. 100 mg of the crushed powder was extracted with 1 ml of extraction solvent (For phytohormone extraction-MS grade methanol/water/formic acid 75:20:5; having 300ng internal standard Adonitol; For pigment extraction-80 percent methanol in MS grade water; having 300ng internal standard Adonitol), by 3-5 min of vigorous vortex, followed by 15 min of sonication. The solution was incubated overnight at −80°C under dark conditions. Next day, the samples were centrifuged at 4°C, maximum speed for 15 min. The supernatant was transferred into a new tube, and reduced to half of its original volume, using centrivap at 4°C, and was filtered through 0.22-micron syringe filter. The LC-MS system (Exion AD, AB Sciex) was used for detection of phytohormones and pigments. Chromatographic separation was done on AB SCIEX Exion LC AD UPLC system, using Phenomenex kinetex core shell C18 column (2.1X150 mm, 1.7µm). Separation of compounds was performed at 0.3 ml/ min flow rate with linear gradient of solvent A (milliQ + 0.1% formic acid) and solvent B (Acetonitrile + 0.1% formic acid) was performed as follows: 0 min 5% B + 95% A; 3 min, 10% B; 6 min, 50% B, 7 min, 70% B; 10 min, 95% B; 12 min, finally column was re-equilibrated to initial ratio of 5% B + 95% A for 4 min with total run time of 16 min. The m/z range for phytohormone and pigment detection was 100-400 and 100-1700, respectively. The sample injection volume was 40 µl for all the samples. Three blank runs were given every time between the biological replicates throughout run. The full scan mass spectrometry data was acquired on the X500R Q-TOF (AB Sciex, USA). The data for all the samples were acquired using electrospray ionization in both positive and negative modes. Mass calibration was performed in positive and negative ion modes before the data acquisition. Data analysis was carried out using the SCIEX OX software provided along with the machine.

### GUS assay

*StREV* promoter transgenic lines (promREV:*uidA*) were grown *in vitro* under LD conditions for 20 d. They were also grown in soil under LD conditions for 8 weeks, followed by SD induction for 15 d. Entire *in vitro* grown plantlets, stolon and tuber samples from SD-induced soil-grown plants were subjected to GUS assay as described in Begum *et al*., (2022) and imaged using olympus MX stereozoom microscope.

### RNA *insitu* hybridization

Longitudinal sections of 8 μm thickness from shoot apex of WT plants were probed with digoxigenin (DIG)-labeled antisense mRNA probes. The probes to detect *StREV* transcripts were PCR-amplified from cDNA using specific primer pairs (Table S5) that amplify part of the cDNA, with T7 RNA polymerase binding site attached to the reverse primer and forward primer. Two independent clones in distinct orientations were used to synthesize the respective sense and anti-sense probes. RNA *insitu* hybridization was carried out as described by Abelenda *et al*., 2019.

### Chlorophyll and anthocyanin estimation

Anthocyanin and chlorophyll estimation from leaves of WT and *miR166* KD lines was performed as per the method in Nakata and Ohme-Takagi (2014) and Bhattacharjee and Sharma (2012), respectively. These experiments were conducted two times, with similar results.

### Histology

For anatomical studies, stems from the middle part of soil-grown plants (6 weeks LD) were used. Free hand transverse sections were cut using a sharp razor blade, and selective sections were stained with Toluidine Blue O solution as described in Pradhan Mitra and Loqué., 2014. The sections were mounted in 70% glycerol, before imaging them on the Nikon Eclipse microscope.

### Dual-luciferase assay

The promoter of *StYUC7* was cloned in the pGreenII0800-LUC vector upstream of the LUC gene. The coding sequence of *5’m-StREV-FLAG* (*StREV* resistant to *miR166*) was cloned in pCB201 downstream to the cauliflower mosaic virus 35S promoter.

Both the constructs were transformed in *A. tumefaciens* strain GV3101. The promoter *StYUC* in pGreenII0800-LUC was co-infiltrated in leaves of *N. benthamiana* with the *5’m-StREV-FLAG* or a negative control empty vector pCB201. Twelve bioreplicates were collected for each condition after 48 hours post-infiltration. Firefly and Renilla luciferase activities were measured using the Dual-Luciferase Reporter Assay System (Promega).

### Yeast-one hybrid assay

The coding sequence of *StREV* and the promoter sequence of *StYUCCA7* (∼3.0 kb upstream) were cloned into the pGEM-T Easy vector (Promega). For preparation of bait expression vector, the promoter was cloned in the destination vector pMW#2 (Addgene) through the restriction sites MluI and SacII. The yeast strain YM4271 was transformed with this bait expression vector (linearised) containing promoter of *StYUC7* and selected in SD-His medium. The prey expression vector was prepared by transferring the coding sequence of *StREV* to the destination vector pDEST-2µ-Gal4-AD via the donor vector pDONR221. The yeast strain Yα1867 was transformed with the prey expression vector and selected in SD-Trp medium. To study the interaction, the prey yeast (Yα1867-StREV) and the bait yeast (YM4271-prom-StYUC7) were mated by mixing in a 1:1 ratio and allowed to grow in YPDA medium. The mated yeast clones were then selected on SD -His, -Trp medium. The interaction was further confirmed by growing the mated yeast clones on SD -His, -Trp medium supplemented with increasing concentrations of 3-amino-1,2,4-triazole (3-AT).

## Data availability

The raw sequencing data were deposited in the NCBI Short Read Archive (SRA) database (http://www.ncbi.nlm.nih.gov/sra/). It can be retrieved using the SRA ID SRP630954. All other relevant data can be found within the manuscript and its supporting materials.

## Acknowledgements

Authors acknowledge Mr. Nilesh Lohakare and Mr. Nitish Lahigude (Indian Institute of Science Education and Research, Pune) for their help in maintaining the plants in the green house facility. We thank Novelgene Technologies Pvt. Ltd. for performing RNA sequencing. We are grateful to Ms. Drishti Kataria (IISER Pune) for her assistance in screening the Y1H clones. We are thankful to Dr. Harpreet Singh Kalsi, Dr. Nilam Malankar, Dr. Kirtikumar Kondhare and Ms. Shikha Singh for their inputs. The authors used ChatGPT (OpenAI) exclusively for language editing and grammar refinement. The scientific content, interpretation, and conclusions are solely those of the authors.

## Funding

NSP and JK are thankful to the University Grants Commission for the Senior Research Fellowship and Junior Research Fellowship, respectively. GA acknowledges funding received from the Department of Biotechnology, India. AKB acknowledges the support and generous funding from the Indian Institute of Science Education and Research (IISER) Pune.

## Authors contributions

N.S.P. designed the experiments, generated transgenic lines, carried out phenotypic and transcriptomic data analyses, conducted experiments and wrote the manuscript draft. A.V. conducted the LC-MS/MS experiments and analyzed its data. B.N. and N.S.P. performed RLM-RACE for target validation. G.A. and N.S.P. generated the *miR166* KD and promREV::uidA transgenic lines. J.K. assisted in molecular cloning and analysis. A.K.B. supervised the study, obtained the funding and resources. N.S.P. and A.K.B. conceived the idea and edited the manuscript. All authors have read and approved the final manuscript.

## Conflict of Interest Statement

The authors declare that they have no conflicts of interest associated with this work.

## Ethics Statement

All experimental research on plants in this study complied with institutional bio-safety guidelines. The plant materials used were maintained in accordance with relevant regulations.

## Supporting Information

**Figure S1.** Details of the precursors and mature *miR166* in potato

**Figure S2.** Abundance of *miR166* in stolons under early LD *vs* SD stages from a photoperiod dependent tuberizing variety ssp. *andigena* 7540

**Figure S3.** Pleiotropic phenotypes of *miR166* suppression on potato

**Figure S4.** PCA and Volcano plots of samples processed for RNA-sequencing

**Figure S5.** Heatmap and RT-qPCR for selective phytohormone genes

**Figure S6.** Heatmap and RT-qPCR for selective tuberization genes

**Figure S7.** Characterization of the *miR166* OE lines

**Figure S8.** Characterization of the REV AS lines

**Figure S9.** GA8 quantification from *in vitro miR166* KD plants

**Figure S10.** Tuber sprout phenotype and heat map for anthocyanin-related genes

**Figure S11.** Development of ectopic sinks in *miR166* KD lines under SD photoperiod

**Figure S12.** Heat maps for selective QTL genes

**Figure S13.** Relative fold change of the *HD-ZIP class III* transcripts

## Supporting Tables

**Table S1.** Differentially expressed genes in WT_Head compared to WT_SSR

**Table S2.** Differentially expressed genes in KD_Head compared to KD_SSR

**Table S3.** Differentially expressed genes in KD_Head compared to WT_Head

**Table S4.** Differentially expressed genes in KD_SSR compared to WT_SSR

**TableS5.** List of Primers

**Table S6.** List of gene IDs

**Supporting Data S1.** MS-MS fragment match for phytohormones and pigments

## Notes

### Competing Interest Statement

The authors have declared no competing interest.

### Summary of Updates

With additional experiments, we now establish that StREV binds to the promoter of StYUCCA7, and upregulates its activity. We have added new results that provide the mechanistic insights of the proposed model. Some textual changes have been made throughout the manuscript.

