## Supplementary Figures 1-13 for "MicroRNA166-REVOLUTA-auxin module affects tuber shape, color and productivity in potato"

### Slide 1
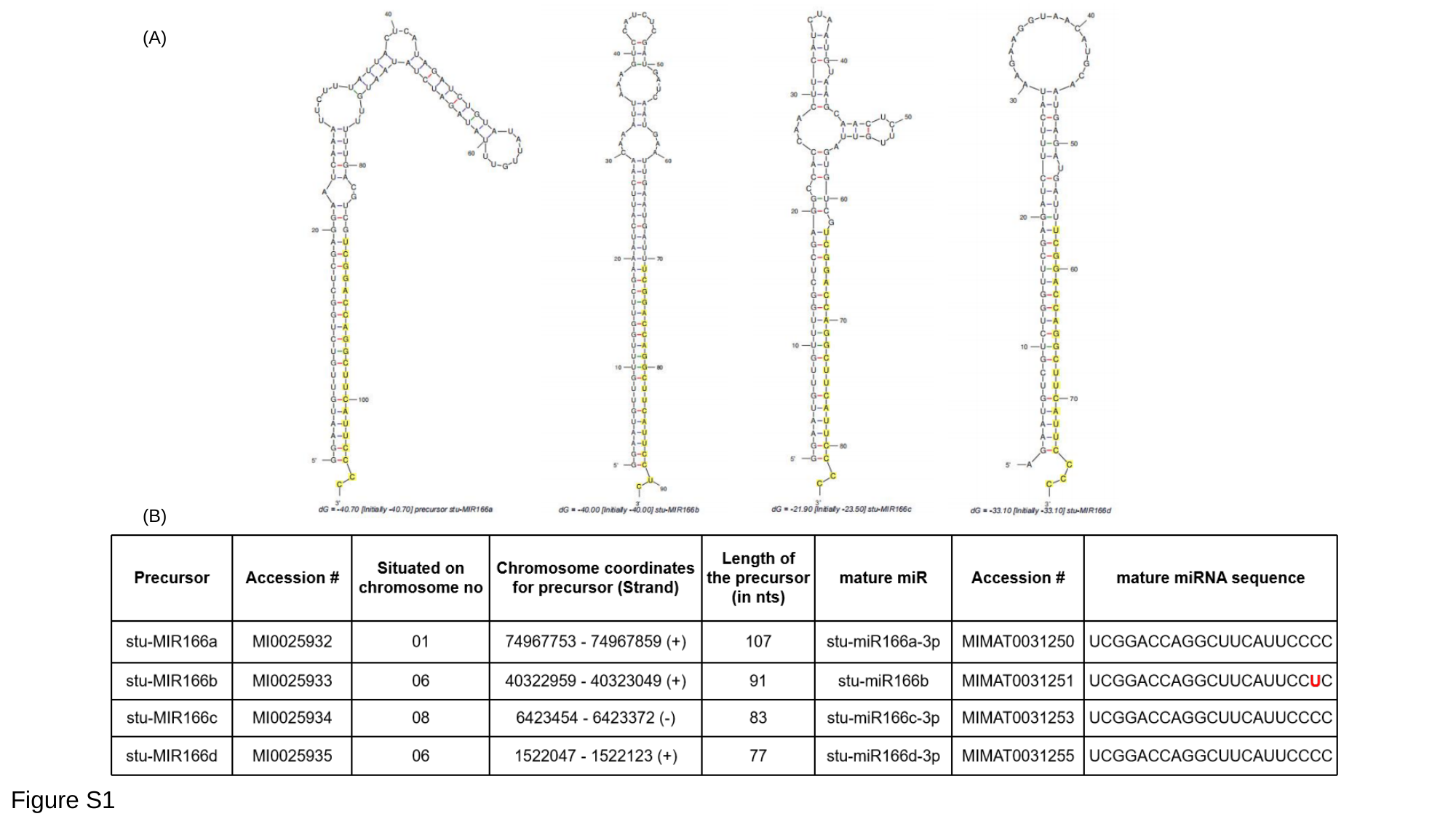

(A)
(B)
Figure S1

### Slide 2
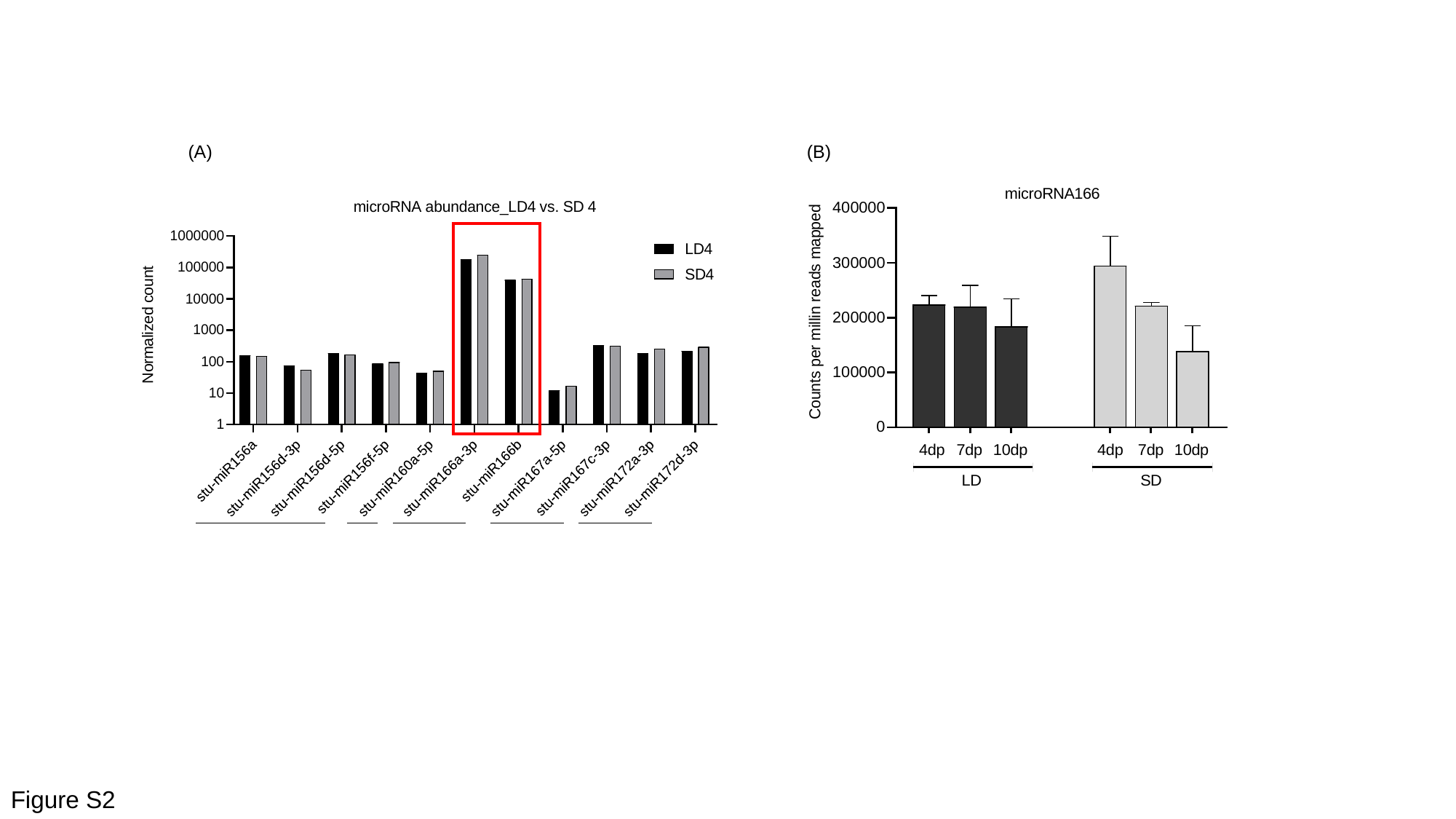

(A)
(B)
Figure S2

### Slide 3
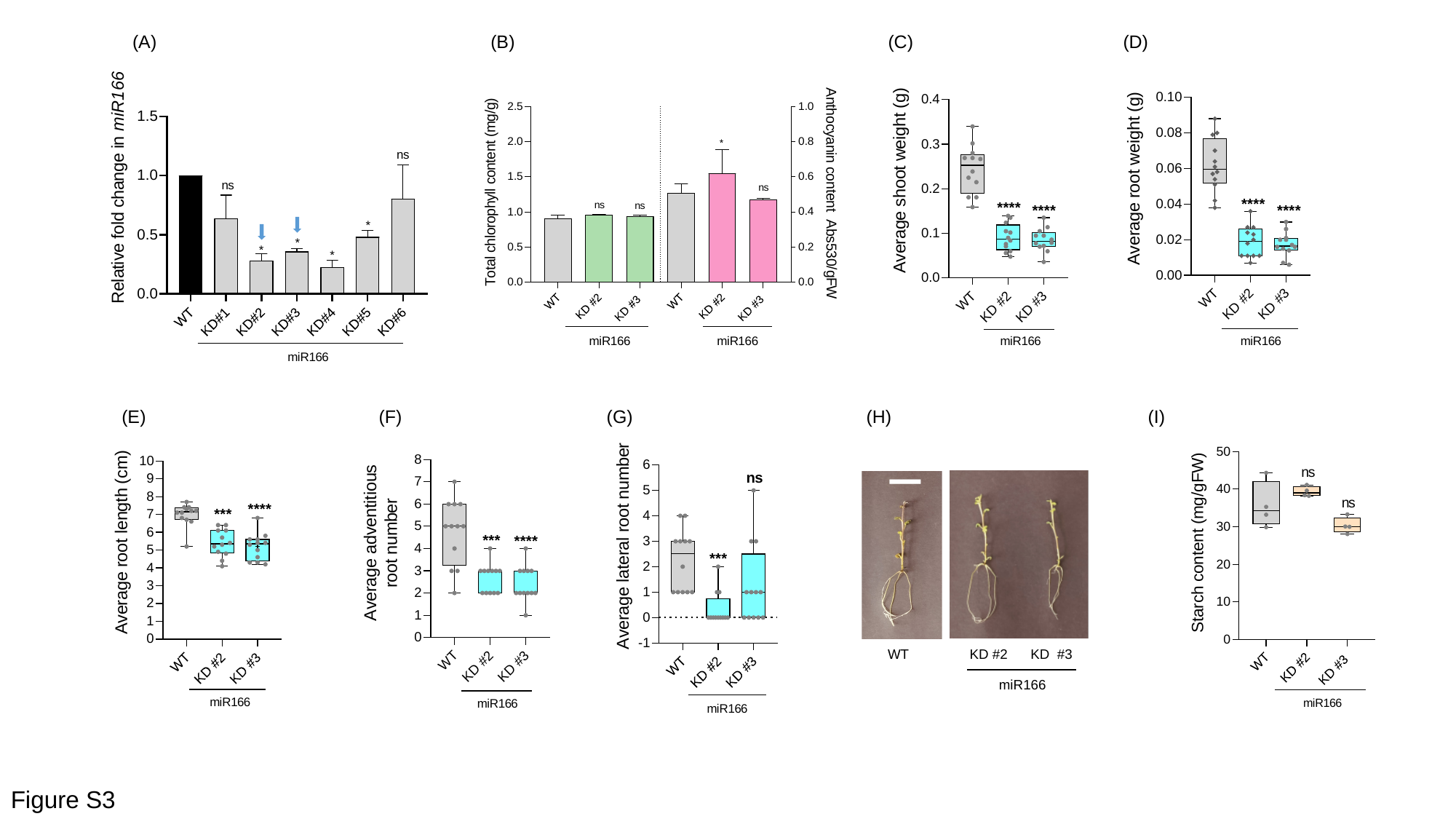

(A)
(B)
(C)
(D)
(E)
(F)
(G)
(H)
(I)
WT KD #2 KD #3
miR166
Figure S3

### Slide 4
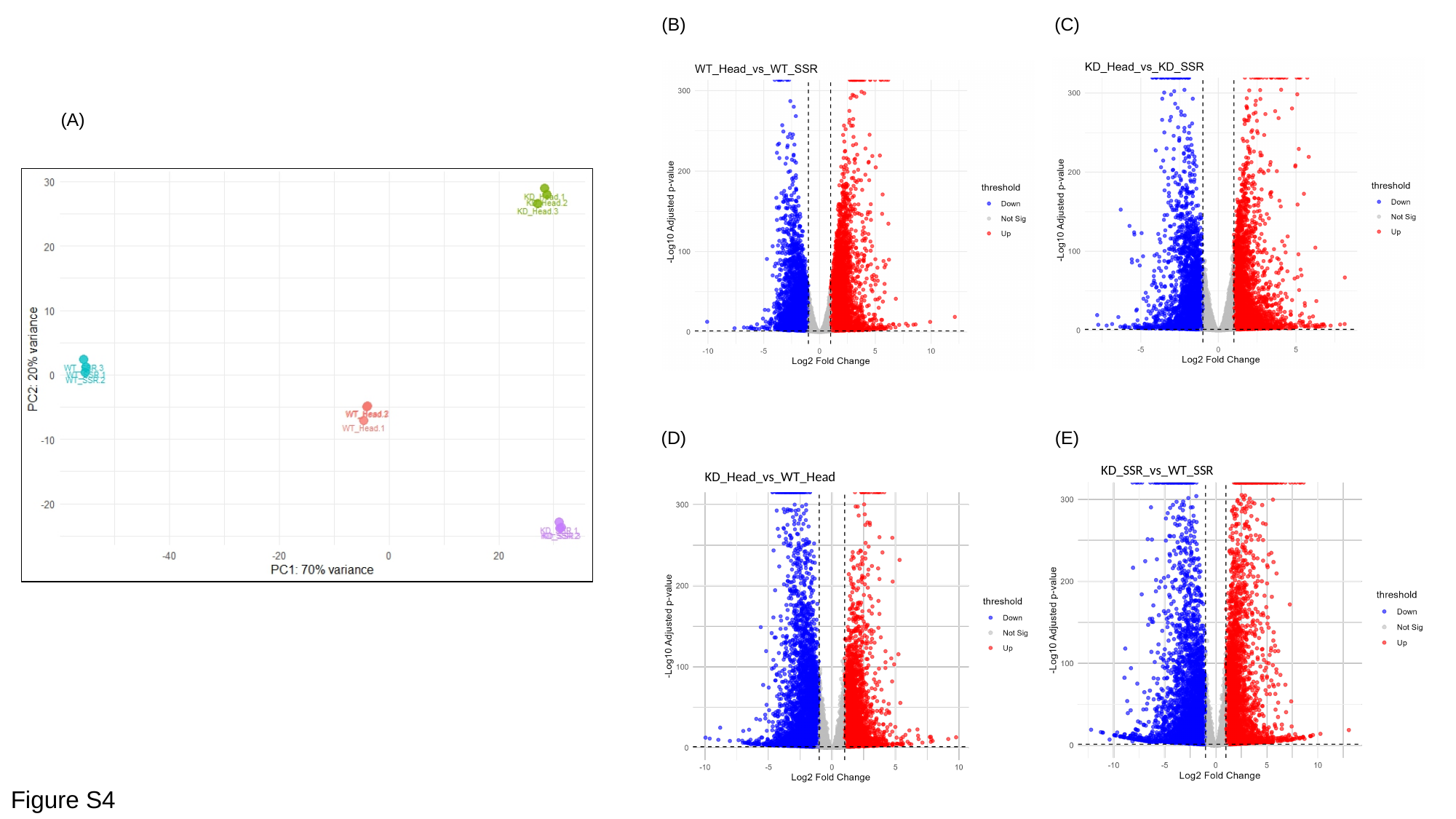

(B)
(C)
(A)
(D)
(E)
KD_SSR_vs_WT_SSR
KD_Head_vs_WT_Head
Figure S4

### Slide 5
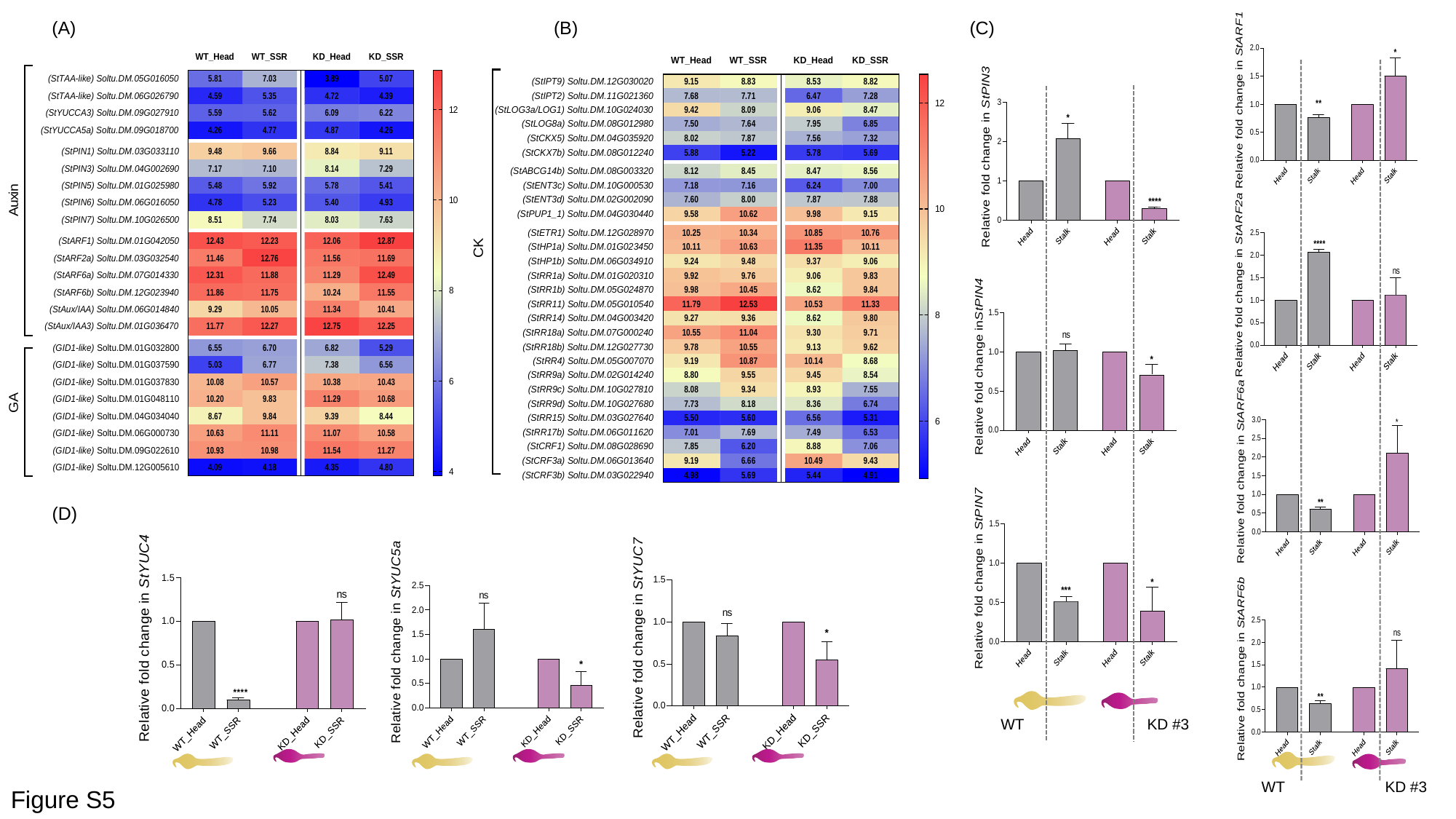

(A)
(B)
(C)
WT
KD #3
(D)
WT
KD #3
Figure S5

### Slide 6
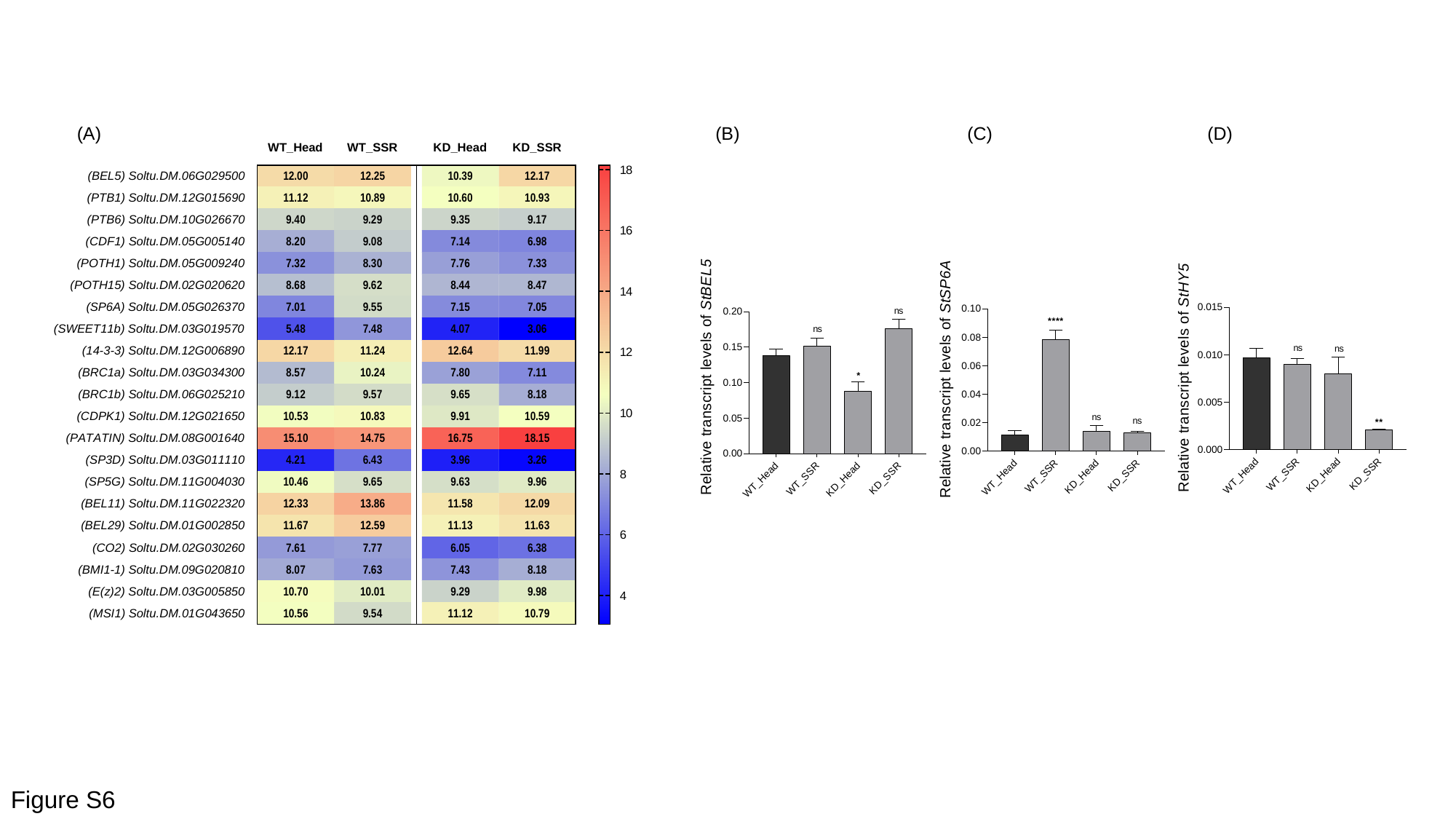

(A)
(B)
(C)
(D)
Figure S6

### Slide 7
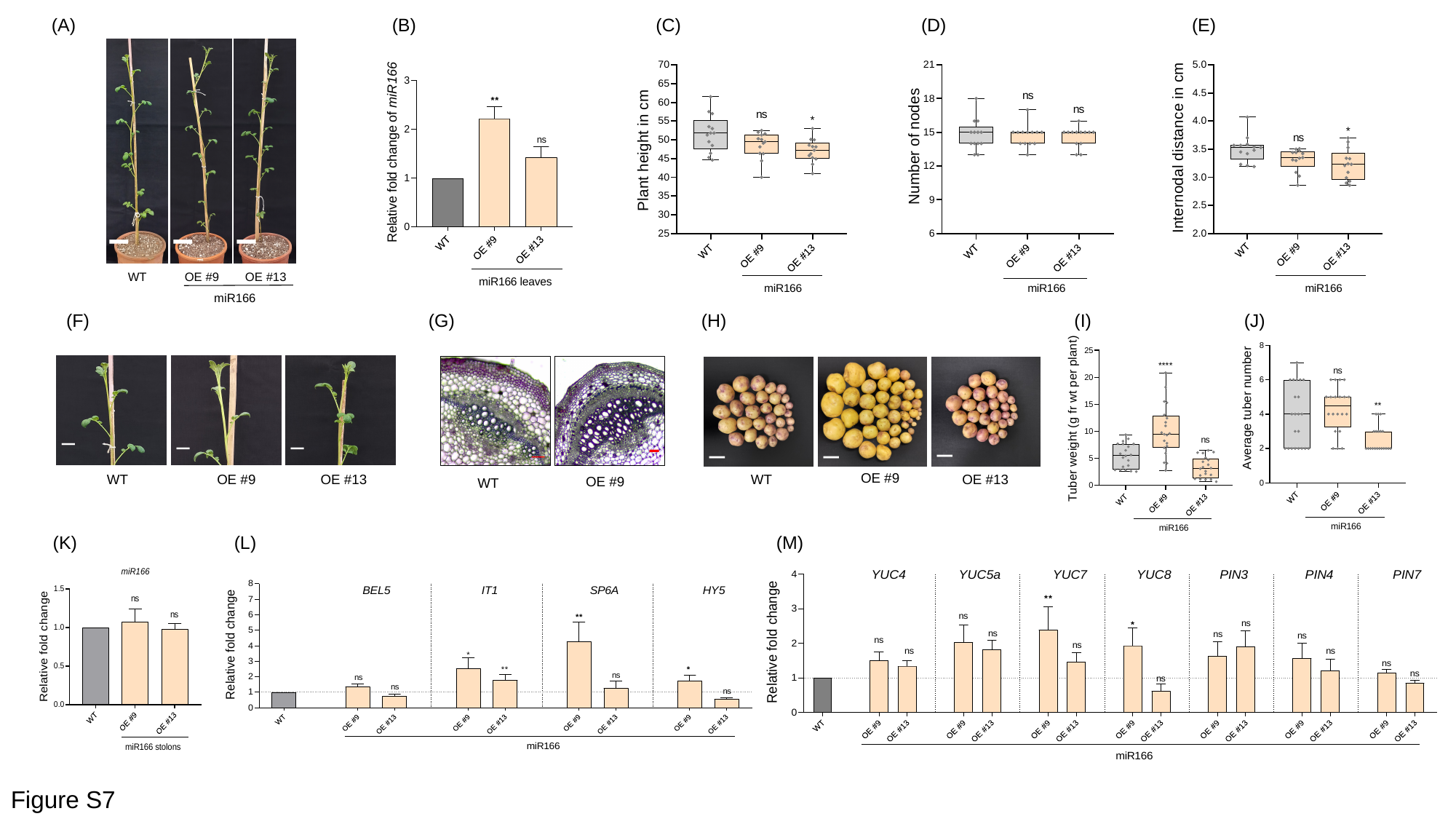

(A)
(B)
(C)
(D)
(E)
WT
OE #9
OE #13
miR166
(F)
(G)
(H)
(I)
(J)
WT
OE #9
OE #13
OE #9
WT
OE #9
WT
OE #13
(K)
(L)
(M)
Figure S7

### Slide 8
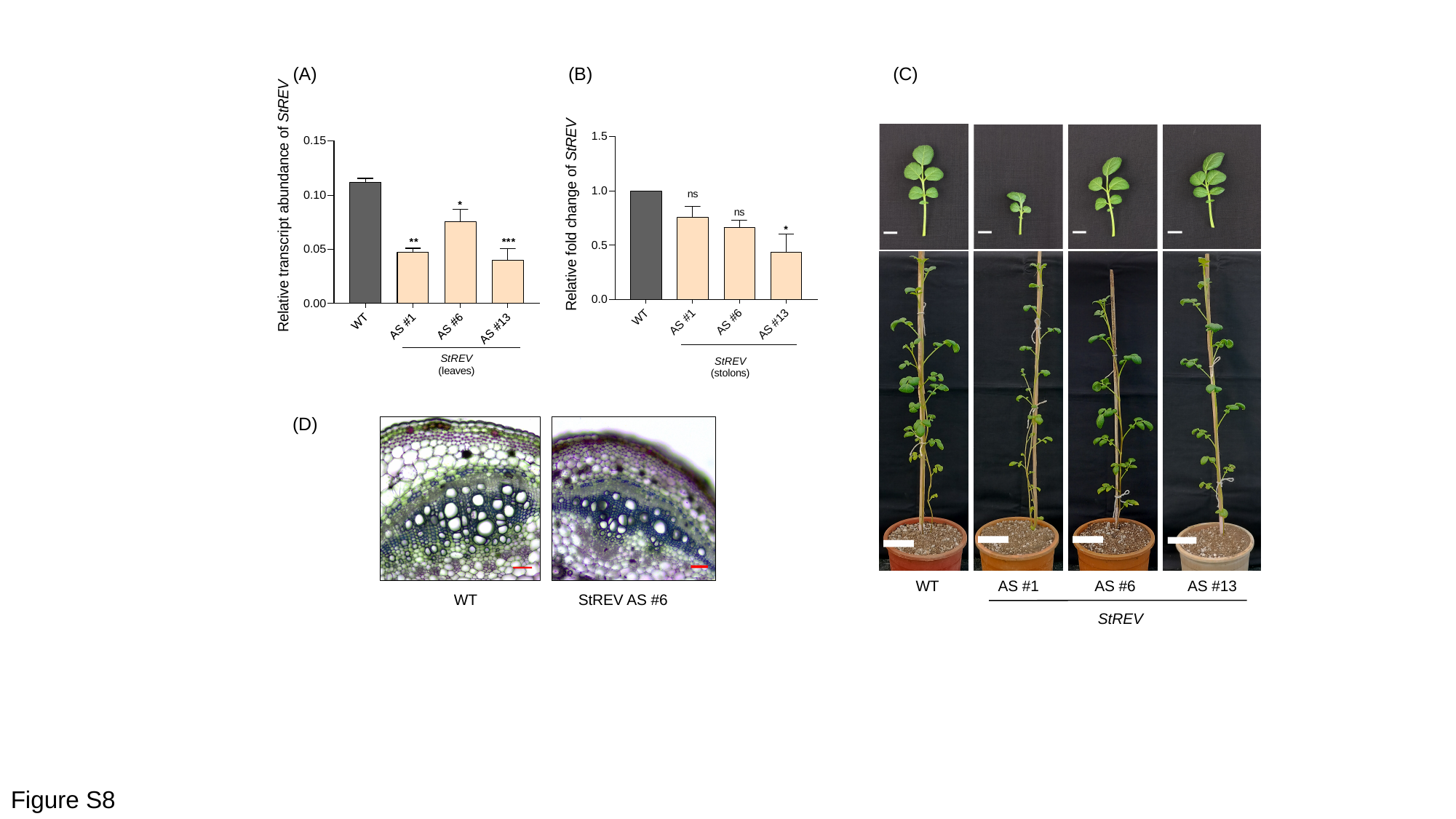

(A)
(B)
(C)
WT
AS #1
AS #6
AS #13
StREV
(D)
WT
StREV AS #6
Figure S8

### Slide 9
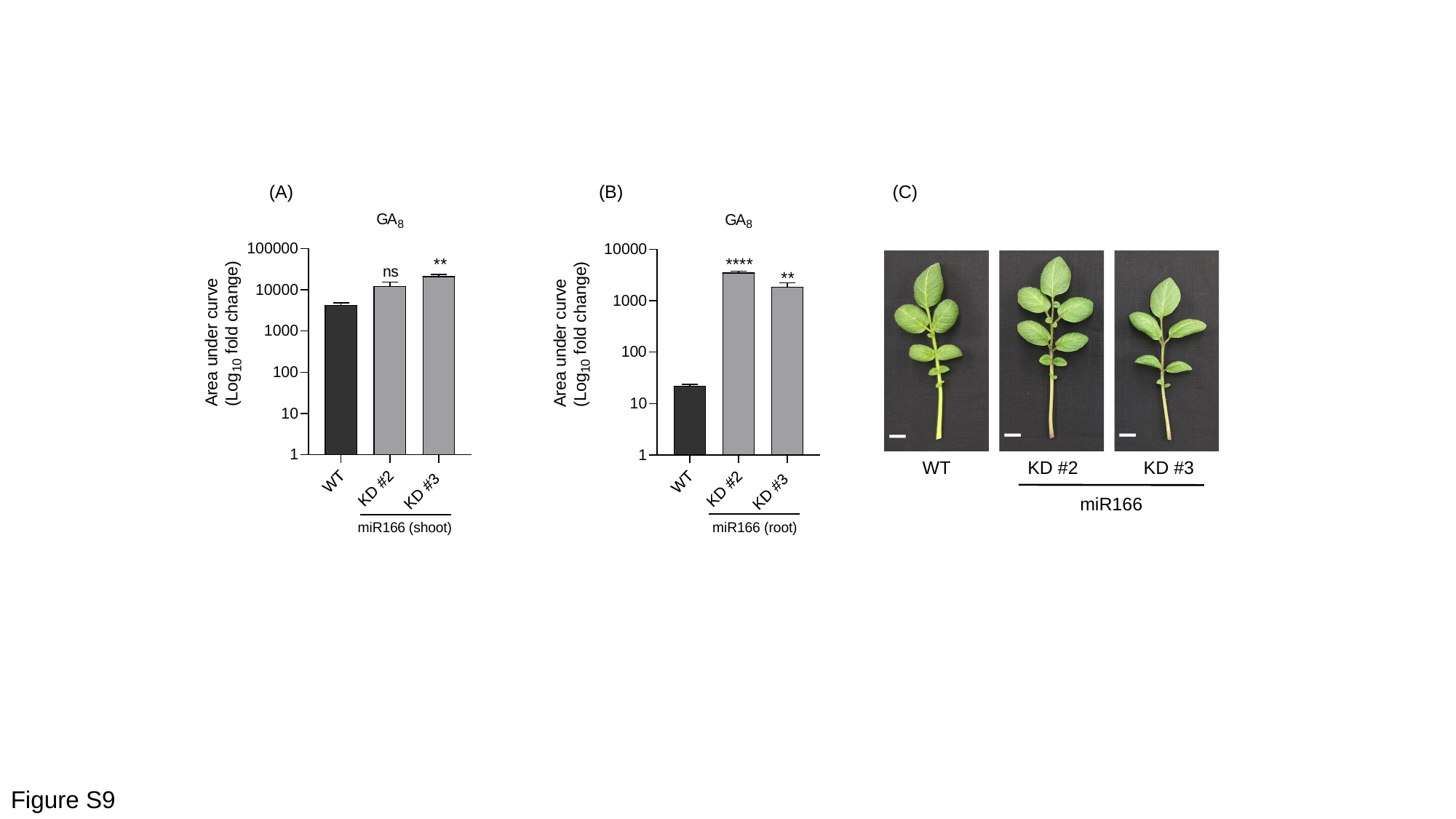

(A)
(B)
(C)
WT
KD #2
KD #3
miR166
Figure S9

### Slide 10
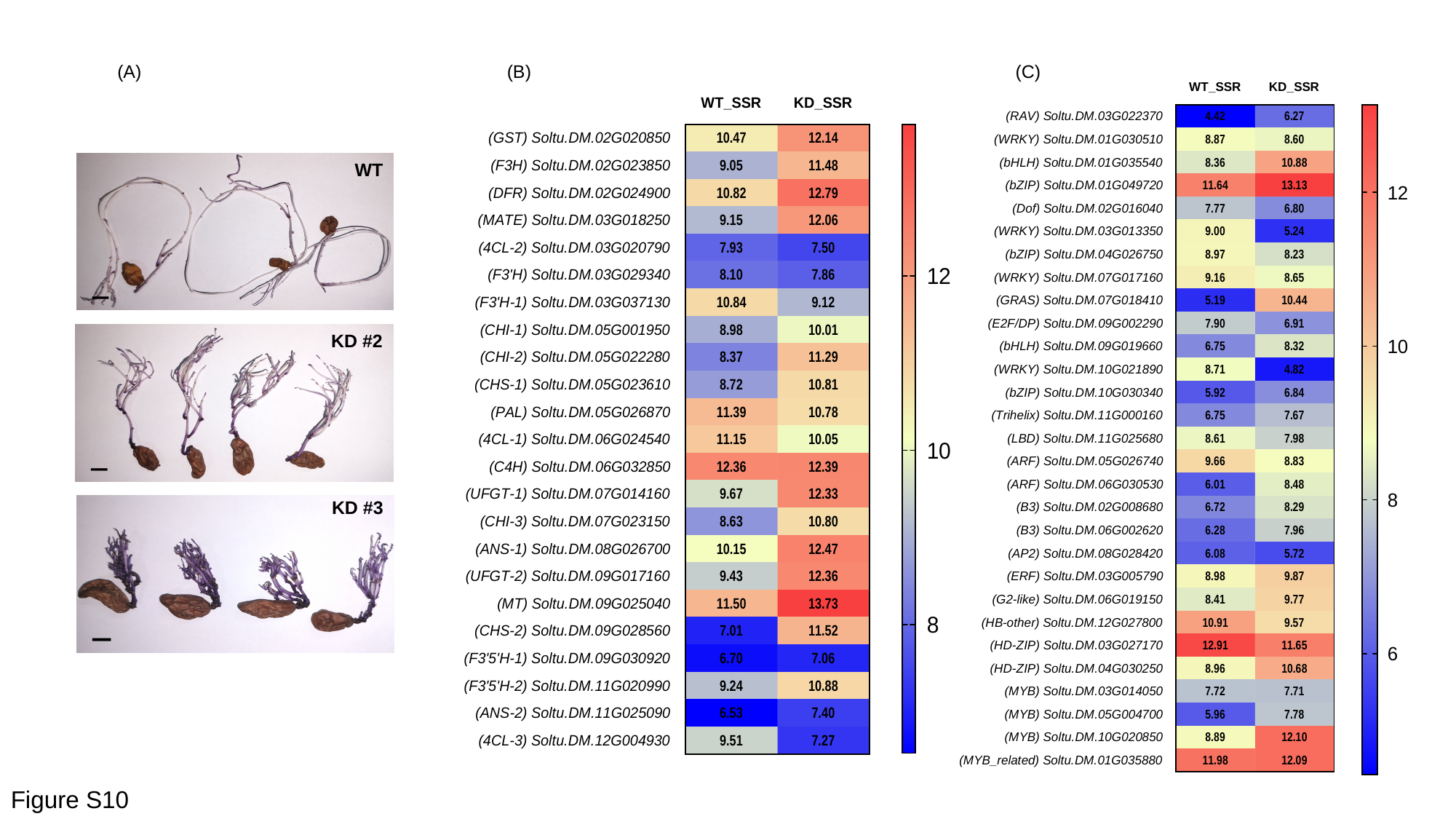

(A)
(B)
(C)
WT
KD #2
KD #3
Figure S10

### Slide 11
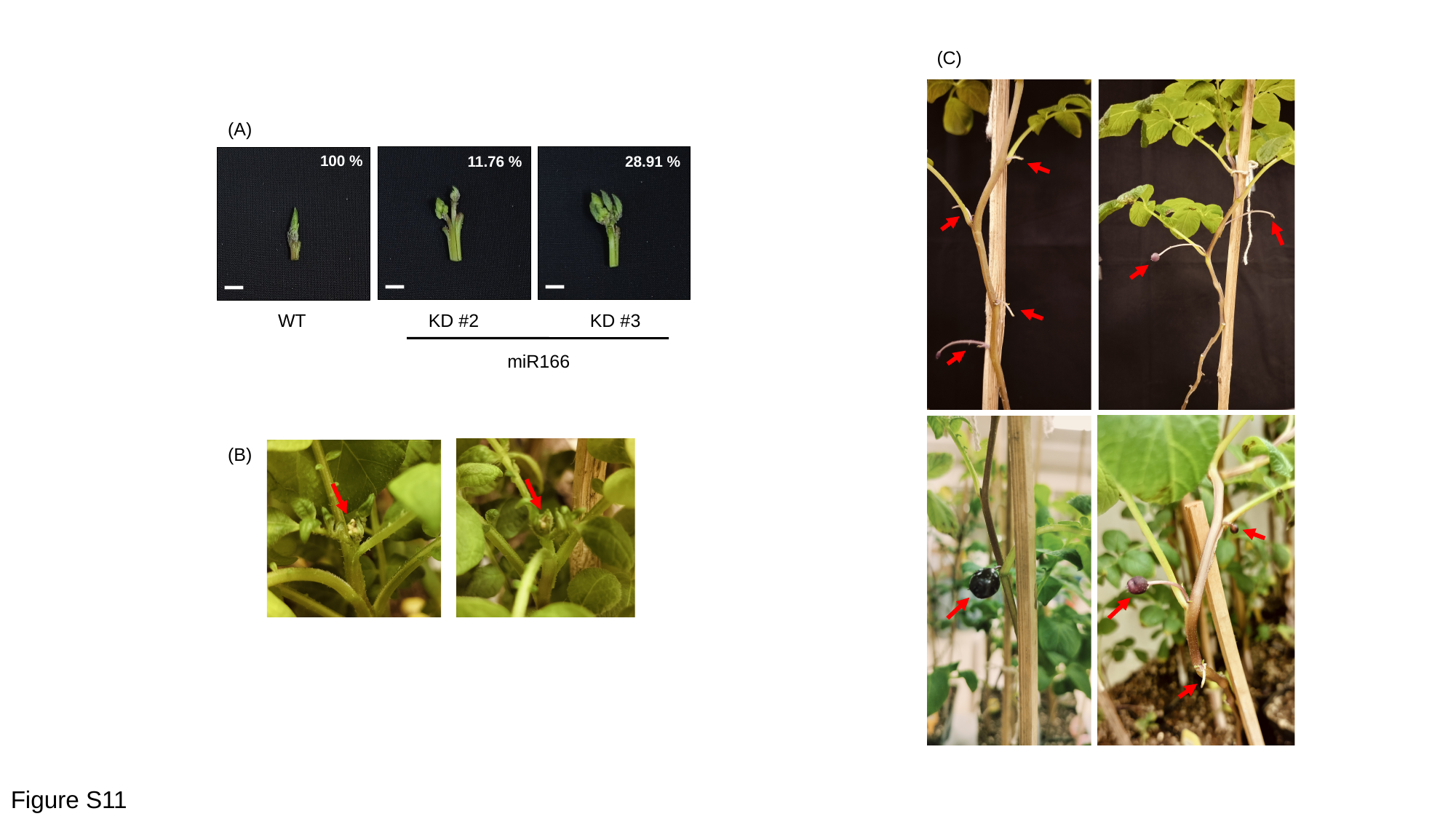

(C)
(A)
100 %
11.76 %
28.91 %
WT
KD #2
KD #3
miR166
(B)
Figure S11

### Slide 12
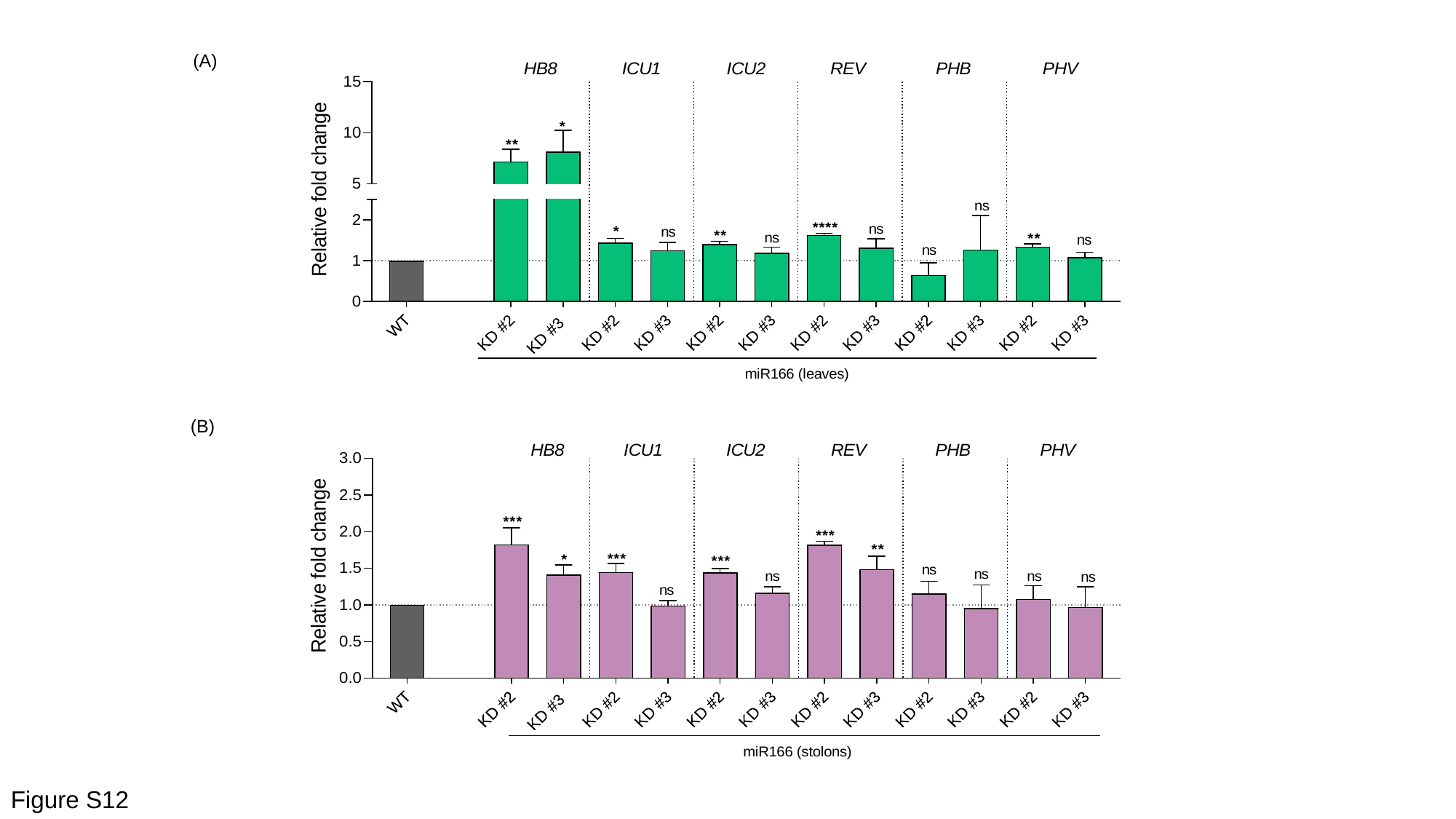

(A)
(B)
Figure S12

### Slide 13
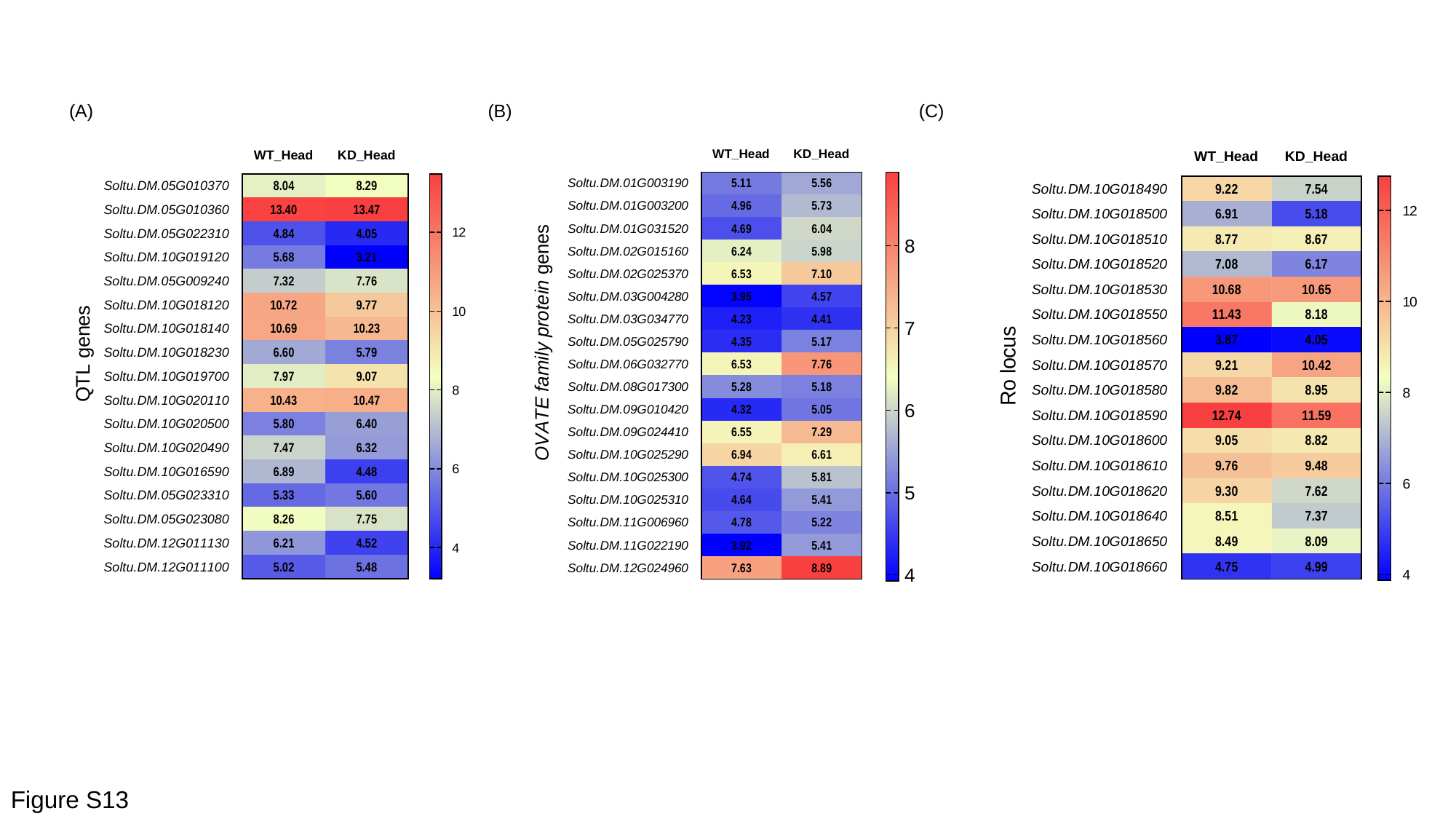

(A)
(B)
(C)
Figure S13
