## Supplementary Information for "MicroRNA166-REVOLUTA-auxin module affects tuber shape, color and productivity in potato"

**Supporting Information**

**Figure S1.** Details of the precursors and mature *miR166* in potato. (A) Secondary structures of *miR166* precursors as predicted by MFold (Zuker, 2003). Mature *miR166* sequences are highlighted in yellow. (B) Details of the precursors and mature *miR166* in potato based on miRBase.

**Figure S2.** Abundance of *miR166* in stolons under early LD vs SD stages from a photoperiod dependent tuberizing variety ssp. *andigena* 7540. (A) *miR166* is the most abundant microRNA amongst all the conserved microRNA families (B) Relative abundance of *miR166* (Counts per million reads mapped; CPM) in stolons under early LD vs SD stages. The small RNA sequencing was done from stolon tissues harvested from photoperiod dependent tuberizing variety ssp. *andigena* 7540 post-photoperiod treatment- 4, 7 and 10 days post-treatment (dp). (A-B) based on small RNA sequencing data from Kondhare *et al.*, 2018.

**Figure S3.** Pleiotropic phenotypes of *miR166* suppression on potato. (A) Relative fold change of *miR166* from the leaves of the stable transgenic lines generated using mimicry construct. Asterisks (one, two, three, and four) indicate significant differences at $p < 0.05$, $p < 0.01$, $p < 0.001$, and $p < 0.0001$, respectively, using Student's t-test. ns = not significant at $p > 0.05$. (B) Total chlorophyll and anthocyanin content from the 5th leaf of *miR166* KD lines and WT plants. *In vitro* root growth assay of *miR166* KD lines: Shoot weight (C), root weight (D), root length (E), adventitious (F) and lateral root numbers (G). (H) Representative images of *in vitro miR166* KD and WT plants. Scale bar = 3cm. (I) Starch content (mg/gFW) of *miR166* KD tubers, relative to WT tubers.

**Figure S4.** PCA and Volcano plots of samples processed for RNA-sequencing. (A) PCA plot for the 12 samples processed for RNA-sequencing. Volcano plots depicting significant DEGs from WT and KD tissues (B-E).

**Figure S5.** Heatmap and RT-qPCR for selective phytohormone genes. (A-B) Heat map was plotted for auxin, GA and CK genes. Numbers inside the box represent variance stabilized transformed (VST) values. (C-D) Validation of selective genes by

RT-qPCR. Students t-test was performed to check significance with one, two, three and four asterisks indicating p-values of < 0.05, < 0.01, < 0.001 and < 0.0001, respectively. ns = not significant. Error bars represent ± SEM from three biological and three technical replicates. *StEIF3e* was used as a reference gene.

**Figure S6.** Heatmap and RT-qPCR for selective tuberization genes. Heat map was plotted for key tuberization pathway genes (A). Numbers inside the box represent variance-stabilized transformed (VST) values. Validation of *StBEL5* (B), *StSP6A* (C) and *StHY5* (D) from the Head and SSR regions of WT and KD tissue. *StEIF3e* was used as reference gene. Error bars indicate standard error of means (SEM) from three biological replicates (n=3). Asterisks (one, two, three, and four) indicate significant differences at $p < 0.05$, $p < 0.01$, $p < 0.001$, and $p < 0.0001$, respectively, using Student's t-test. ns = not significant at $p > 0.05$.

**Figure S7.** Characterization of the *miR166* OE lines. (A) Plant phenotype of the *miR166* OE lines, compared to the WT plants. Scale bar = 5 cm. (B) Relative fold change of *miR166* in leaves of *miR166* OE lines. Plant height (C), number of nodes (D) and internodal distance (E) from *miR166* OE lines in comparison with WT (n=13). The leaf curvature (F) did not seem to change in the OE plants. Scale bar = 1 cm. (G) Transverse sections of stem from *miR166*OE line along with WT tissue. Scale bar = 50 microns. (H) Representative images for tubers from *miR166* OE lines compared to WT, each image contains tubers from 10 plants per line. Scale bar = 2 cm. Tuber weight (I) and average tuber number (J) for *miR166*-OE lines compared to WT. Data was taken from soil grown plants after a two months of SD induction, and plotted from 20 individual plants per line (mean ±SEM). Relative fold change of *miR166* (K), few tuber marker genes (L) and auxin biosynthesis and transport genes (M) from the stolons (7 days SD) of *miR166* OE. *StEIF3e* was used as reference gene. Error bars indicate standard error of means (SEM) from three biological replicates (n=3). Asterisks (one, two, three, and four) indicate significant differences at $p < 0.05$, $p < 0.01$, $p < 0.001$, and $p < 0.0001$, respectively, using Student's t-test. ns = not significant at $p > 0.05$.

**Figure S8.** Characterization of the REV AS lines. Relative levels of *REV* transcripts from leaves (A) and stolons (B) of REV AS lines. *StEIF3e* was used as reference gene.

Error bars indicate standard error of means (SEM) from three biological replicates (n=3). Asterisks (one, two, three, and four) indicate significant differences at $p < 0.05$, $p < 0.01$, $p < 0.001$, and $p < 0.0001$, respectively, using Student's t-test. ns = not significant at $p > 0.05$. (C) Representative plant images of REV-AS lines, compared to the WT.(D) Transverse section of REV AS stem. Scale bar= 50 microns.

**Figure S9.** GA8 quantification from *in vitro miR166* KD plants. GA8 quantification from *in vitro* shoots (A) and roots (B) of *miR166* KD plants (C) Representative image of mature leaflets (5th leaflet from Apex) from the *miR166* KD lines, compared to the WT.

**Figure S10.** Tuber sprout phenotype and heat map for anthocyanin-related genes (A) Tuber sprouts of *miR166* KD lines showed increased branching. The tuber sprouts from *miR166* KD lines are stunted, profusely branched and pigmented, compared to WT sprouts. The image is taken after 40 weeks of tuber harvest. Scale bar = 1 cm. Heat map depicting up regulation of several genes encoding enzymes in the anthocyanin pathway (B) and some of the TF families having the strong correlation with these anthocyanin pathway genes (C) (based on Zhang et al., 2024) from the *miR166* KD_SSR tissue. Numbers inside the box represent variance stabilized transformed (VST) values.

**Figure S11.** Development of ectopic sinks in *miR166* KD lines under SD photoperiod. Images of floral buds, excised (A) (Scale bar = 0.5 cm) and intact (B) from soil grown MIM166 plants. Numbers in top right corner in (A) denote the percentage of plants having vegetative apex in WT, and floral buds in KD #2 and KD #3. *miR166* KD plants developed aerial stolons/tubers under SD photoperiod (C).

**Figure S12.** Heat maps for selective QTL genes. Heatmap for QTL genes (A), Ro locus (B) and Ovate family protein genes (C) from WT and *miR166* KD tissues. Values inside the box represent VST normalized gene expression count.

**Figure S13.** Relative fold change of the *HD-ZIP class III* transcripts. Relative fold change of *HD-ZIP class III* in the young leaves (A) and stolons (B) of soil grown plants, compared to the WT. *StEIF3e* was used as reference gene. Error bars indicate standard error of means (SEM) from three biological replicates (n=3). Asterisks (one, two, three, and four) indicate significant differences at $p < 0.05$, $p < 0.01$, $p < 0.001$, and $p < 0.0001$, respectively, using Student's t-test. ns = not significant at $p > 0.05$.

**Supporting Tables**

**Table S1.** Differentially expressed genes in WT_Head compared to WT_SSR

**Table S2.** Differentially expressed genes in KD_Head compared to KD_SSR

**Table S3.** Differentially expressed genes in KD_Head compared to WT_Head

**Table S4.** Differentially expressed genes in KD_SSR compared to WT_SSR

**TableS5.** List of Primers

**Table S6.** List of gene IDs

**Supporting Data S1.** MS-MS fragment match for phytohormones and pigments
