## Supplementary Data S1 for "MicroRNA166-REVOLUTA-auxin module affects tuber shape, color and productivity in potato"

MS-MS fragments of phytohormones were obtained from stolon-to-tuber transition stages and in vitro plant tissues using variable collision energy. The experimentally obtained fragments were used to match with fragments generated by CFM-ID computationally and standard phytohormones. We identified IAA-3, $GA_3$ and $GA_8$, and their MS-MS fragments match is provided below.

(A) Standard IAA-3

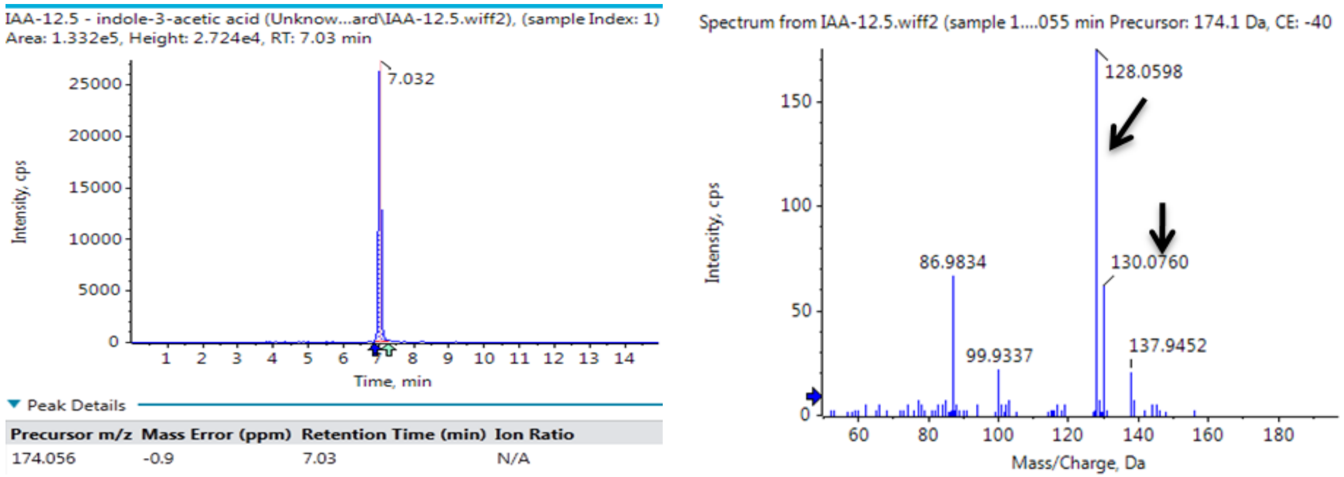


(B) IAA-3 from samples

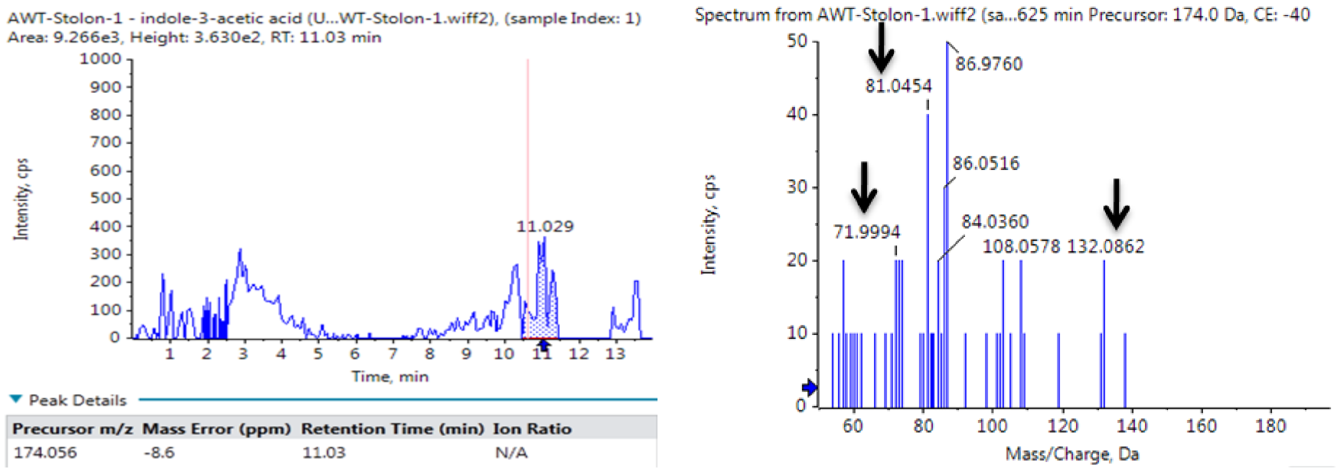

(C) Standard Gibberellic Acid 3

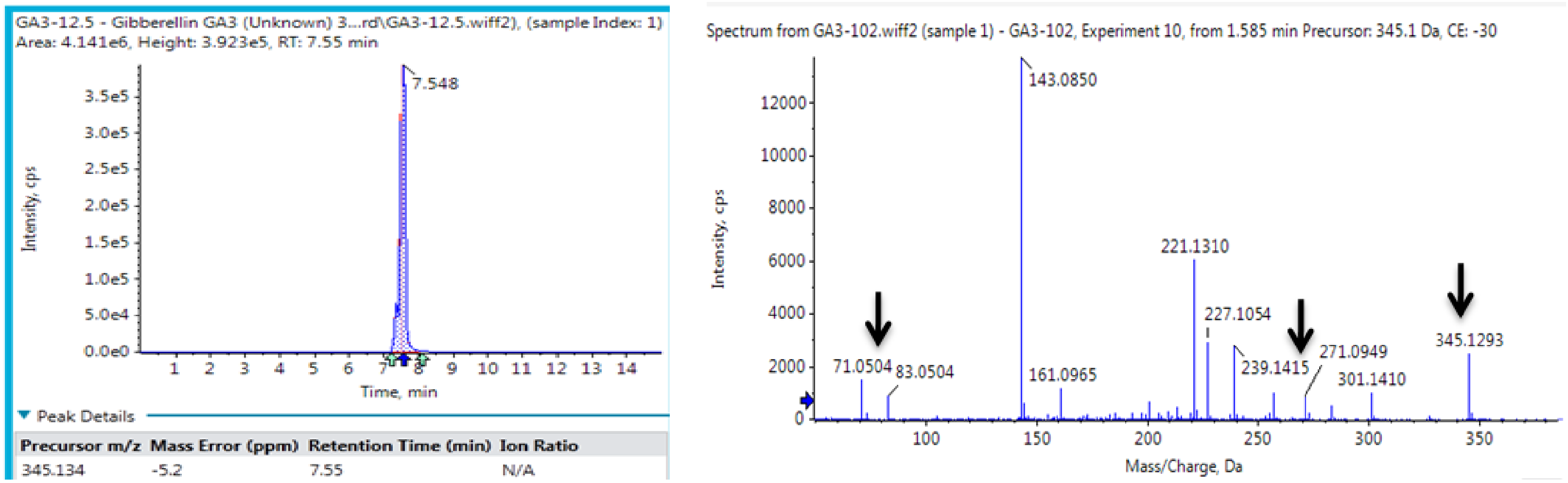


(D) Gibberellic Acid 3 from samples

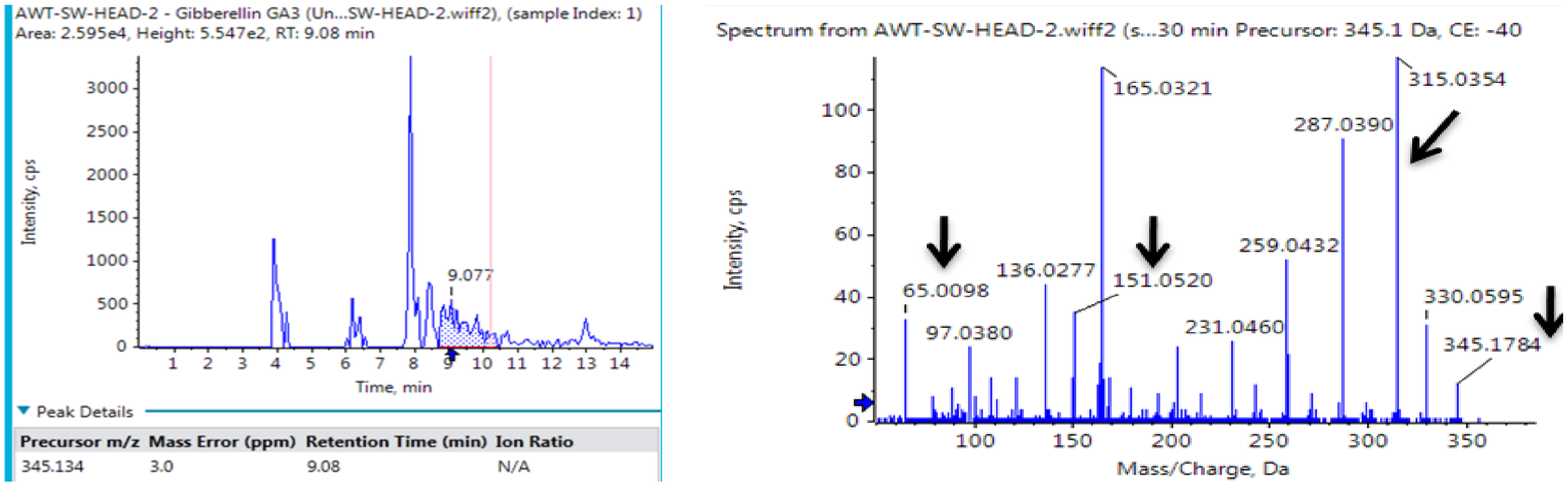

(E) GA 8 from samples

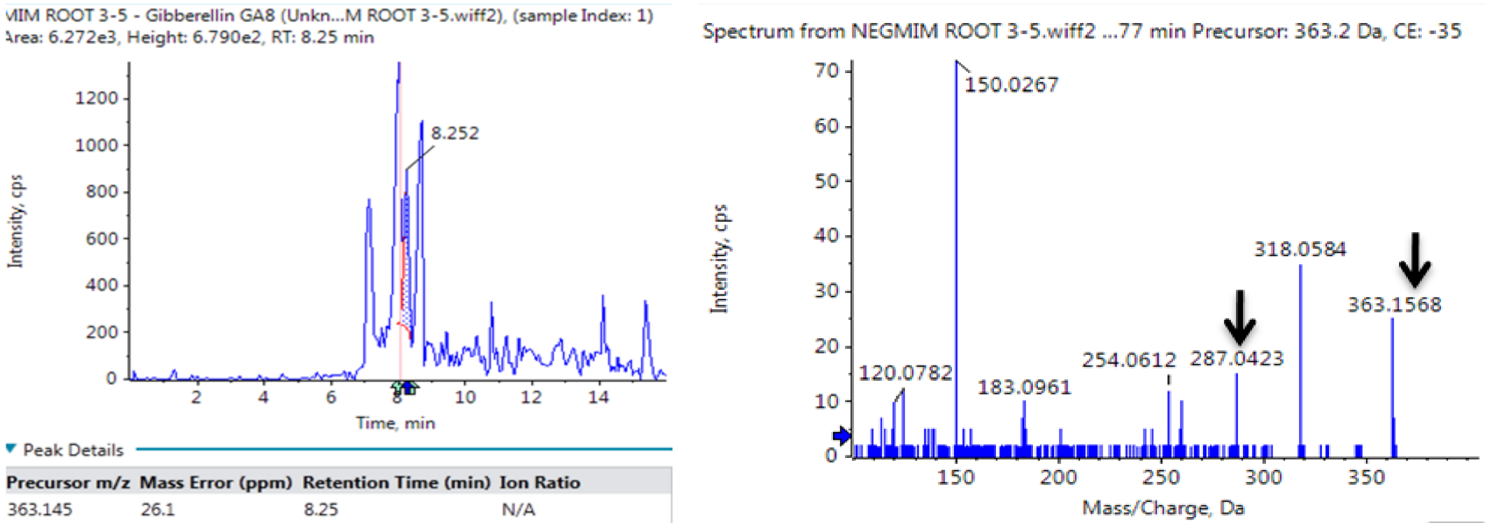

MS-MS fragments of anthocyanins were obtained from peel tissues using variable collision energy. The experimentally obtained fragments were used to match with the fragments generated by CFM-ID computationally. Their MS-MS fragments data is provided below.

(A) Cyanidin-3-glucoside

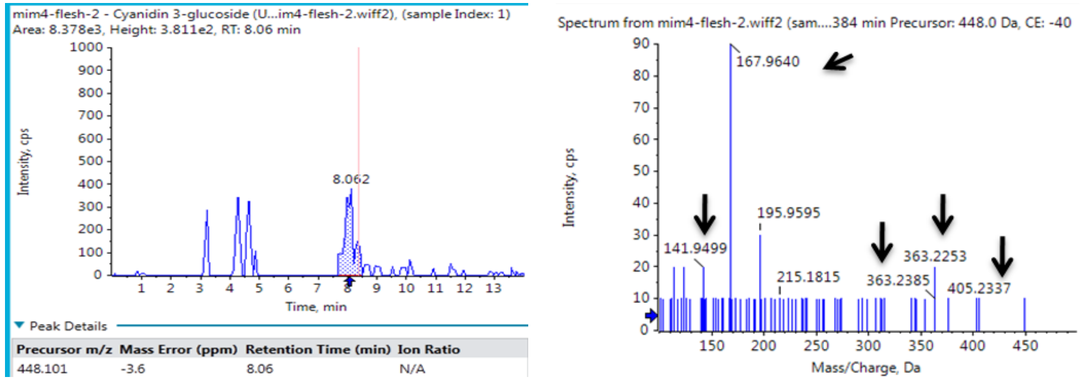


(B) Delphinidin-3-glucoside

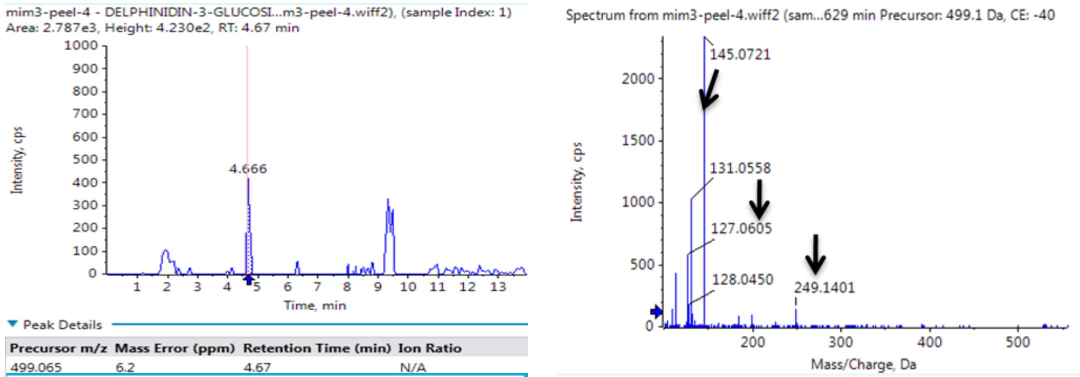


(C) Pelargonidin-3-glucoside

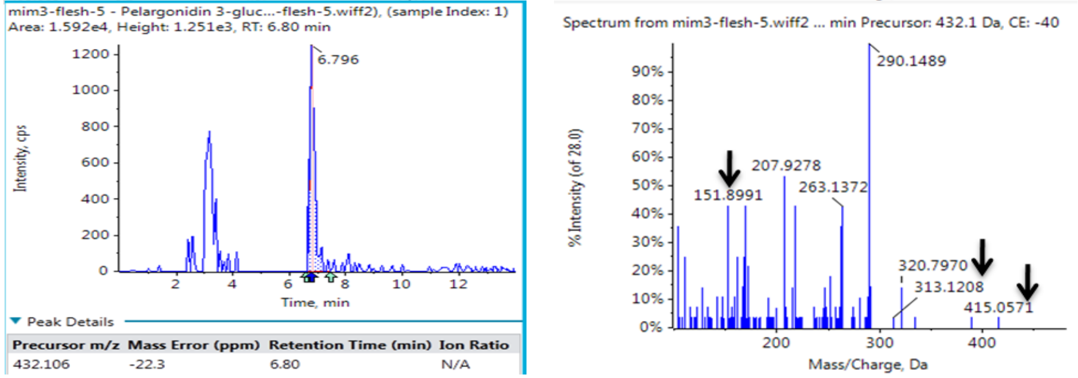
